# Zebrafish larval nitrogen excretion is flexible and resilient to loss of rhesus glycoproteins

**DOI:** 10.64898/2026.08.28.747819

**Authors:** Wouter Mes, Robin Haanen, Arslan Arshad, Peter H.M. Klaren, Marcel J.M. Schaaf, Erin Faught, Tsutomu Nakada, Maartje A.H.J. van Kessel, Marnix Gorissen

**Author notes:** These authors contributed equally (Maartje A.H.J. van Kessel, Marnix Gorissen).

## Abstract

Nitrogenous waste excretion is essential for all developmental stages of fish. Embryonic fish excrete urea, transitioning to cutaneous and later branchial ammonia excretion. In zebrafish, ammonia excretion involves rhesus glycoproteins Rhbg and Rhcgb in keratinocytes and ionocytes, but the developmental moment they appear in the gill remains unclear. Potential redundancy between Rhbg and Rhcgb in ammonia excretion is also not fully investigated, nor is the difference in response to low pH. We hypothesized that rhesus glycoproteins are partially redundant, and that they differ in their response to low pH as ammonia excretion enables ionocytes to exchange Na^+^ and H^+^ (Rh-NHE-metabolon). We predicted that a loss of *rhbg* or *rhcgb* induces compensatory responses. We characterized the transition from urea to branchial ammonia excretion from 0 to 8 days-post fertilization (dpf) and the response to pH 5.0 on the expression and localization of rhesus glycoproteins in control zebrafish and *rhbg* or *rhcgb*-crispants. Effects of high external ammonia (HEA, 500 µM NH_4_Cl) and 10 mM HEPES-buffering were further characterized in *rhcgb*-crispants. Rhag and Rhbg appeared in the gill at 5 dpf, while Rhcgb appeared at 6 dpf. A loss of *rhbg* or *rhcgb* did not impact baseline N-excretion, illustrating that zebrafish can maintain ammonia excretion without the full complement of rhesus glycoproteins. We observed no compensatory increase in rhesus glycoproteins, but expression of the transporter *hippocampus-abundant transcript 1b* increased. HEA-exposed *rhcgb-*crispants switched to urea as primary nitrogen waste. Together, these findings underline the plasticity of the larval in dealing with nitrogenous waste.

**Summary statement:** We targeted zebrafish ammonia transporters *rhbg* and *rhcgb* with CRISPR-Cas9 mutagenesis to characterize larval nitrogen excretion and found high resilience to disruptions of ammonia excretion, including a switch to urea excretion.

## Introduction

As adults, most teleost fish are ammonotelic, which means that the majority of their produced nitrogenous waste is directly excreted as ammonia into the environment, and not detoxified to other nitrogen species (Ip and Chew, 2010). Gills are the principal organ for ammonia excretion in adult fish, which is one of its key functions together with gas exchange, osmoregulation, and acid-base regulation (Evans et al., 2005). In contrast to adult teleost fish, embryonic teleosts are ureotelic (*i.e.,* they excrete urea rather than ammonia as principal nitrogenous waste) before hatching. The energy-costly detoxification of ammonia to urea (costing 5 moles of ATP per 1 mole of urea) in pre-hatch fish embryos is necessary to prevent ammonia accumulation, as they are surrounded by a small volume of chorionic fluid, a chorion and an unstirred boundary layer around the egg that prevent ammonia diffusion(Bucking et al., 2013; Zimmer et al., 2017). After hatching, zebrafish switch from ureotely to ammonotely and excrete ammonia via the skin, as their gills are not yet fully developed and their surface-to-volume ratio is large enough for cutaneous excretion to suffice (Zimmer et al., 2017).

In both cutaneous and branchial ammonia excretion mechanisms, rhesus glycoproteins play an important role. There are multiple rhesus glycoproteins which have been shown to transport ammonia in both marine and freshwater fish (Nakada et al., 2007a; Nakada et al., 2007b). Zebrafish (*Danio rerio* (Hamilton, 1822)) have five rhesus glycoprotein genes (*rhag, rhbg, rhcga* [previously named *rhcg2*]*, rhcgb* [previously named *rhcg1*] and *rhcgl1* [previously named *rhcg3*]) (Nakada et al., 2007a). The mechanism of ammonia excretion and its relationship to other gill physiological functions (*e.g*., Na^+^-uptake, acid-base balance) are well-defined in zebrafish (Shih et al., 2008; Shih et al., 2013): Rhbg and Rhcgb have been shown to transport ammonia in zebrafish and are expressed in different cells in both larval and adult stages, presenting two separate ammonia excretion pathways. Rhbg is present in skin keratinocytes in 3-4 day old larvae (Shih et al., 2013) and can be found on gill pavement cells in adult fish (Braun et al., 2009b), while Rhcgb expression is found in a subtype of ionocytes known as H^+^-ATPase-rich (HR) ionocytes in larval skin and adult gill tissue (Nakada et al., 2007a).

HR-ionocyte-expressed Rhcgb is thought to play a key role beyond ammonia excretion, in Na^+^-uptake via Na^+^/H^+^-exchanger 3b (NHE3b), as it indirectly provides the outward H^+^-gradient necessary to take up Na^+^ against its own gradient (Wright and Wood, 2009; Ito et al., 2013). Since this process is even more challenging at low pH (due to a higher external H^+^-concentration), reliance on ammonia excretion as acid equivalent and counter-ion may indeed be even greater in these circumstances. Indeed, several studies showed increased ammonia flux at a lower pH (Shih et al., 2008; Kumai and Perry, 2011; Wright et al., 2016). The role of ammonia as nitrogenous waste, but also as acid equivalent and counter-ion in Na^+^-uptake, makes it highly relevant and challenging to assess the role of different rhesus glycoproteins in nitrogen physiology (Zimmer, 2024).

While the contribution of the aforementioned rhesus glycoproteins to ammonia excretion is well-attested, the relative contribution of separate rhesus glycoproteins and alternative ammonia transporters to the overall process is less clear. For example, morpholino knockdown of *rhag*, *rhbg* and *rhcgb* all reduced ammonia excretion (Shih et al., 2008; Braun et al., 2009a) but *rhcgb*-knockout fish exhibited *increased* ammonia excretion (Zimmer and Perry, 2020), possibly through compensation via alternate rhesus glycoproteins. Additionally, the importance of each rhesus glycoprotein may differ between developmental stages: at 4 days post-fertilization (dpf) these proteins mostly localize to the skin (Braun et al., 2009a) but by 8 dpf, whole-mount *in situ* hybridizations show expression of *rhag, rhbg* and *rhcgb* in the gills. Whether this change in localization affects relative contribution to nitrogen excretion and whether there are differences between rhesus glycoproteins in the timing of this change is currently unknown. Recently discovered hippocampal-abundant transcript (Hiat) proteins 1a and 1b (*hiat1a, hiat1b*) ammonia-transporting proteins may provide zebrafish with further flexibility in dealing with ammonia excretion (Zhouyao et al., 2022).

Understanding how the complex set of ammonia transporters help zebrafish cope with environmental stressors relating to nitrogen excretion is highly relevant: zebrafish may serve as model to assess the risk of environmental acidification and nitrogenous waste pollution on freshwater fish and echanistic insights into the complex process of nitrogen excretion may provide us with tools to mitigate impacts of global environmental change and understand how fish can adapt to these adverse conditions (Zimmer et al., 2021). Exposure to high external ammonia (HEA) inhibits ammonia excretion at first, but induces gene expression of rhesus glycoproteins to re-establish an outward ammonia gradient (Braun et al., 2009b). Removal of the acid-trapping mechanism through buffering of the embryo medium also reduces ammonia flux (Shih et al., 2008; Kumai and Perry, 2011).

In this study, we characterized the role of rhesus glycoproteins in zebrafish during the larval stage and their contributions to the overall process of nitrogen excretion. We first confirmed the expression and localization of rhesus glycoproteins in the skin and gill by analyzing whole body transcript levels and using immunohistochemistry (IHC). Rhag and Rhbg appear in epithelial gills cells at 5 dpf, whereas ionocyte-expressed Rhcgb localizes to the gill from 6 dpf. To examine the role of both pathways in ammonia excretion, we subjected the larvae to low environmental pH (5.0) and employed an *rhbg*- and *rhcgb*-crispant knockdown approach (Kroll et al. (2021). Importantly, we observed no changes in nitrogen excretion in either crispant and no compensation via rhesus glycoproteins, but we did observe marked upregulation of Hiat1a/b in *rhcgb-*crispants. We hypothesized that exposure of *rhcgb-*crispants to high environment ammonia (HEA) levels or increased buffering would elucidate the mechanisms that underpin the resilience of larval zebrafish to disturbances in nitrogen metabolism. In response to HEA, *rhcgb*-crispants excreted most nitrogen as urea, suggesting that ammonia excretion is no longer feasible or energetically favorable and showing that loss of rhesus glycoproteins can be overcome through alternate ammonia transporters or metabolic adaptations.

## Methods

### Zebrafish husbandry

Animal experiments were approved by the Dutch Central Authority for Scientific Procedures on Animals (license number AVD1030020198606). Adult zebrafish (AB strain) were housed in groups of 20 to 30 fish in 4-L tanks in a laboratory Recirculating Aquaculture System (pH 7.7-8.2; 27°C; 300 µS cm^-1^; 14 h light, 10 h dark). The water was recirculated through a separate fixed-bed biofilter and UV-treated after biofiltration. The fish were fed twice per day with Gemma micro 300 (3% w/w Body Mass; Skretting Nutreco N.V., Amersfoort, the Netherlands) and once per day with live *Artemia* sp. In order to obtain zebrafish embryos, fertilization was performed by natural spawning at the beginning of the light period. Eggs were collected, washed, placed in Petri dishes with 20 ml E3 medium (5 mM NaCl, 0.17 mM KCl, 0.33 mM CaCl_2_, 0.33 mM MgSO_4_, 0.0001% methylene blue (Brand et al., 2002)), and kept in a 28.5°C incubator (14h light, 10 h dark).

### Experimental series

#### Characterization of nitrogen excretion mechanisms and rhesus glycoprotein localization

To study the development of nitrogen excretion mechanisms of zebrafish larvae, we reared wild-type zebrafish embryos from fertilization until 8 days post-fertilization (dpf) and measured nitrogenous waste excretion every day. At 6 hpf, unfertilized eggs were removed and phenylthiourea (PTU; final concentration of 0.004%) was added to E3 medium in order to prevent the development of pigmentation in the embryos used for immunohistochemistry. Dead embryos or larvae were removed daily and E3 or E3+PTU medium was refreshed at 2 dpf, 5 dpf, and 7 dpf.

#### Nitrogenous waste excretion measurements

For ammonia and urea determination, embryos or larvae were transferred to 2 mL tap water from the fish facility. Water samples (1 mL) were taken at 0 and 3 hours, and immediately frozen. Ammonium concentrations were measured using the orthophthaldialdehyde (OPA) method (Taylor et al., 1974), with the reagent volume modified for use in a 96-well plate (200 µL OPA reagent and 10 µL sample). Urea determination in water samples was performed by adding 90 µl urease suspension (6 mg mL^-1^ in dH_2_O) to 90 µL of sample. After incubation at 30°C for 20 min, ammonium was measured as described above and the urea concentration in the sample was calculated.

#### Sampling

Every day (1-8 dpf), zebrafish from the same clutch as those used for nitrogenous waste excretion (*n* = 60) were sampled to analyze expression of nitrogen metabolism genes and ionocyte markers. Zebrafish larvae were euthanized using an overdose of 2-phenoxyethanol (0.1% v/v, pH 7.5) and pooled for RNA isolation and analysis (*n* = 6; depending on developmental stage, 5 fish/sample for 1-2 dpf, 4 fish/sample for 3-5 dpf and 3 fish/sample for 6-8 dpf). Zebrafish were collected in 2-mL Eppendorf tubes and E3 medium was removed, after which they were snap-frozen in liquid nitrogen and stored at -80°C until RNA isolation.

For IHC analysis, zebrafish larvae were treated with the H^+^-ATPase-rich (HR) ionocyte vital stain concanavalin A (ConA) conjugated to a FITC fluorescent probe before sampling (Li et al., 1995). Zebrafish larvae (3, 5, 6 and 7 dpf, *n* = 60 per developmental stage) were transferred to a Petri dish containing E3 medium with 50 µg mL^-1^ ConA-FITC and incubated in the dark for 30 min. After ConA treatment, animals were briefly rinsed in E3 medium and then euthanized with an overdose of 2-phenoxyethanol (0.1% v/v, pH 7.5). The samples were then fixed overnight in 4% paraformaldehyde in Phosphate Buffered Saline (1x PBS – 137 mM NaCl, 10 mM Na_2_HPO_4_, 2.7 mM KCl, 1 mM KH_2_PO_4_) at 4°C, washed three times in PBS-T (1x PBS, 0.1% v/v Tween-20) for 5 min and stored in 100% methanol at -20°C until IHC analysis.

#### Mechanistic contribution of rhesus glycoproteins during low environmental pH

To study the effects of environmental pH on zebrafish nitrogen metabolism and the localization of rhesus glycoproteins, wildtype zebrafish embryos were reared from fertilization until 5 dpf in E3 medium at either pH 5.0 (low pH) or pH 7.4 (control). At 6 hpf, unfertilized eggs were removed and PTU (final concentration of 0.004%) was added to E3 medium in order to prevent the development of pigmentation in the embryos used for IHC. Dead embryos or larvae were removed daily and E3 or E3+PTU medium was refreshed at 1 dpf and 3 dpf.

#### Sampling

On 4 and 5 dpf, zebrafish larvae were sampled to measure expression of nitrogen metabolism genes and ionocyte markers. Zebrafish larvae were euthanized using an overdose of 2-phenoxyethanol (0.1% v/v, pH 7.5) and pooled for RNA isolation and analysis (*n* = 6; 4 fish/sample). Zebrafish were collected in 2-mL Eppendorf tubes and E3 medium was removed with a pipette, after which they were snap-frozen in liquid nitrogen and stored at -80°C until RNA isolation. For IHC analysis, zebrafish larvae were treated with the H^+^-ATPase-rich (HR) ionocyte vital stain, ConA conjugated to a FITC fluorescent probe before sampling, similar to described above. After ConA treatment, animals were briefly washed in E3 medium and then euthanized with an overdose of 2-phenoxyethanol (0.1% v/v, pH 7.5). The zebrafish were fixed overnight in 4% paraformaldehyde in 1xPBS at 4°C, washed three times in PBS-T for 5 min and stored in 100% methanol at -20°C until immunohistochemical analysis.

#### Generation of rhbg- and rhcgb-crispant (F0) larvae

To understand the role of rhbg- and rhcgb-mediated ammonia excretion in zebrafish larvae, we used CRISPR-Cas9 mutagenesis to create F0 crispant zebrafish that have biallelic knockouts of either *rhbg* or *rhcgb* gene. This was done by injecting three guides targeting different exons of each gene, a strategy previously described in detail for zebrafish (Kroll et al., 2021). Ribonucleoprotein (RNP) complexes (guide RNA + Cas9 protein) were generated in a similar way as described in Kroll et al. (2021): CRISPR RNA (crRNA) for the targeted exons was designed using ChopChop (https://chopchop.cbu.uib.no/) (*rhbg* exon 1: GCGATAGTAGAACTCATTCGTGG; exon 2: CACCATGGCAAGATCCACGTTGG; exon 4: TCCATGACTATTCATACATTTGG, *rhcgb* exon 1: GTTTGTGCGGTATAACGAGGAGG; exon 2: CGCAGCTTTCGGCCTCCAGTGGG; exon 3: TGGGATCACGCTGTATGCTGTGG). Guide RNA (gRNA) was produced by annealing the crRNA and universal tracrRNA (IDT, Leuven, Belgium) at 95°C (5 min). RNPs were then assembled by hybridizing the guide RNA to Cas9 protein (IDT, Leuven, Belgium) at 37°C for 5 min. These RNPs were then injected into fertilized wild-type embryos prior to the single cell stage. The efficiency of mutagenesis was evaluated by PCR using primers targeting each affected exon of *rhbg* and *rhcgb* (Table S1). This was performed on 8 F0-crispants per group and efficiency was assessed by measuring shifts in melt curve temperature of the amplicons as described in Faught and Schaaf (2025) The efficiency was above 90% in all cases. Furthermore, absence of the protein was confirmed in each larva used for IHC. To control for the effects of RNP injection itself, a third group of zebrafish was injected with scrambled (nonsense) guide RNA linked to the Cas9 protein.

#### Nitrogenous waste excretion of rhbg- and rhcgb-crispant zebrafish and sampling

*Rhbg*- and *rhcgb-*crispants (F0) as well as control-injected larvae were then reared under the same conditions as described to 5 dpf. At 5 dpf, we measured nitrogenous waste production of pooled groups of larvae (*n* = 10) during 3 hours (as described in experiment 1) and sampled larvae from the same batch for gene expression analysis (*n* = 9-15 per condition, 3 larvae per sample) and immunohistochemistry (*n* = 12 per condition).

#### Functional relevance of rhesus glycoproteins during ammonia-related challenges

For *rhcgb-*crispants and control-injected larvae, we subsequently tested the response to high external ammonia (HEA) and strongly buffered medium (10 mM HEPES, pH 7.4). For each condition, we placed groups of 10 larvae in 2-mL Eppendorf tubes (*n* = 2-4 per condition) containing 1 mL E3 medium (control), 1 mL E3 medium with 500 µM NH_4_Cl (HEA), or 1 mL E3 medium with 10 mM HEPES (HEPES-exposure) and measured the accumulation of nitrogenous waste during 3 hours. Following this measurement, the medium was removed and zebrafish were snap-frozen in liquid nitrogen and stored at -80°C until RNA isolation.

#### RNA isolation & real-time quantitative PCR

Total RNA was extracted from zebrafish embryos and larvae using TRIzol (Invitrogen, Carlsbad, CA, USA) reagent according to manufacturer’s protocol with an adjusted TRIzol volume (400 µL) and a second precipitation in sodium acetate and ethanol. After removal of residual genomic DNA by treatment with RNAse-free DNase (BioRad, Hercules, CA, USA), 500 ng of RNA was reverse transcribed with iScript reverse transcriptase (BioRad, Hercules, CA, USA) in a 20-µL mixture containing iScript Reaction Mix and RNAse I inhibitor for 5 min at 25°C, 20 min at 46°C and 1 min at 96°C. After reverse transcription, cDNA samples were diluted with 140 µL ultra-pure H_2_O and 4 µL were used for qPCR reactions.

Relative expression of rhesus glycoproteins (*rhag, rhbg*, *rhcga*, *rhcgb*, and *rhgcl1*), ionocytes markers Na^+^/K^+^-ATPase, H^+^-ATPase, Na^+^/H^+^-exchanger 3b (*atp1a1a, atp6v1aa, nhe3b*) and urea metabolism genes (*cpsIII, otc, ass1, asl1*, and *ut*) was measured with real-time quantitative PCR (RT-qPCR). In short, 4 μL of the cDNA was used as template in a reaction with 10 µL SYBR Green Master Mix (Biorad, Hercules, CA, USA), 0.4 µM forward and 0.4 µM reverse primers (Table 1) and ultrapure H_2_O in a total reaction volume of 20 µL. RT-qPCR was carried out using a CFX 96 qPCR machine (Biorad, Hercules, CA, USA) with the following conditions: 3 min at 95°C, followed by 40 cycles of 15 s at 95°C and 1 min at 60°C. All expression data were normalized to an index of two reference genes (*elongation factor 1a* (*elf1a*) and *40S ribosomal protein S11* (*rps11*)) (Vandesompele et al., 2002).

**Table 1:**
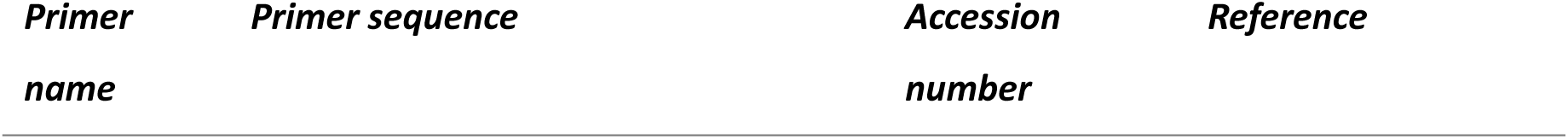
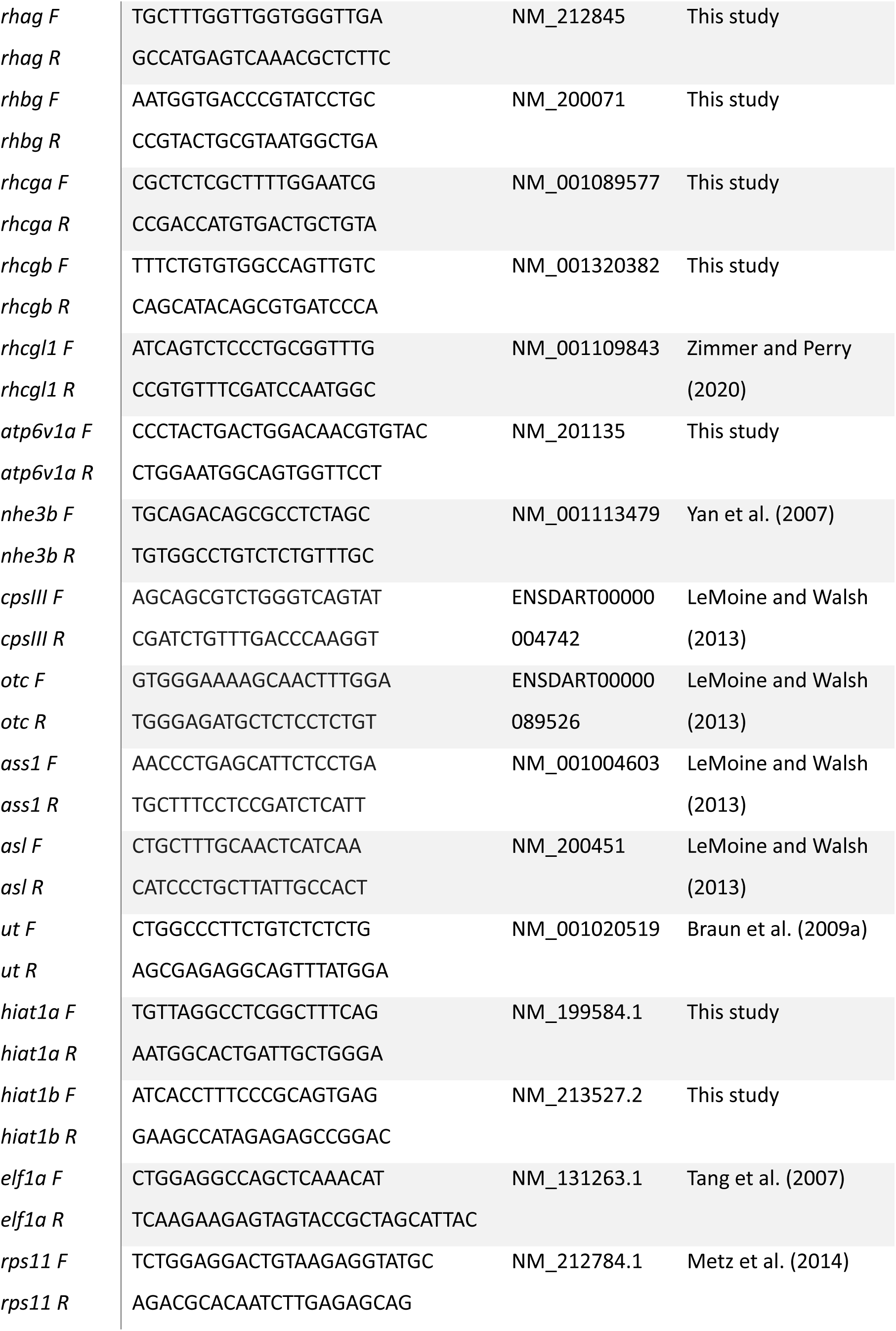
Primer sequences used for gene expression studies of developing zebrafish

#### Immunohistochemistry (IHC)

Fixed and dehydrated larvae (3-7 dpf) were rehydrated stepwise to PBS-T (75% - 50% - 25% methanol in PBS-T) and rinsed four times in PBS-T. Larvae were then permeabilized by incubation in 10 µg mL^-1^ proteinase K in PSB-T at room temperature (RT) for 30 min. Subsequently, the larvae were rinsed for 5 min in PBS-T and fixed for 20 min in 4% paraformaldehyde in PBS and rinsed three times in PBS-T. Following fixation, larvae were incubated in blocking buffer (10% normal donkey serum [NDS] in PBS-T) for 3 hours at RT. Afterwards, larvae were incubated overnight at 4°C with primary antibodies under agitation. Four antibodies against different rhesus glycoproteins were generously shared by Prof. S. Hirose (Tokyo Institute of Technology): fRhag, fRhbg and fRhcg2, polyclonal rabbit antibodies developed against pufferfish (*Takifugu rubripes*) rhesus glycoproteins (Nakada et al., 2007b) and zebrafish (d)Rhcg1, a polyclonal rabbit antibody developed against the Rhcgb (previously Rhcg1) protein of zebrafish (Nakada et al., 2007a). Rhag and Rhcg1 were used in 1:500 dilutions (in 1% NDS in PBS-T), Rhbg and Rhcg2 were used in 1:250 dilutions (in 1% NDS in PBS-T). After primary antibody incubation, larvae were washed four times in PBS-T under agitation. Then, larvae were incubated with a fluorescent secondary antibody (donkey anti-rabbit IgG conjugated with Cy3) at 1:500 dilution (in 1% NDS in PBS-T) for 2 hours at room temperature under agitation. From this step onwards, samples were shielded from light with aluminum foil to protect the fluorescent label. Larvae were washed three times in PBS-T and subsequently transferred to 50% glycerol in 1xPBS and stored at -20°C until microscopic analysis. Images were taken using a confocal laser scanning microscope (Leica SP8X, Mannheim, Germany) and image processing was performed in Fiji (Schindelin et al., 2012).

#### Statistics

Statistical analyses were carried out using Graphpad Prism version 10.2 for Mac (Graphpad software, La Jolla, USA). Testing for normal distribution of data was done using the Shapiro-Wilk test. If data were normally distributed, parametric statistical tests were used for further analyses. In case of a single factor, we tested whether standard deviations of the groups were equal (tested for using an F-test), and if so, a regular one-way analysis of variance (ANOVA) was used with Dunnett’s multiple comparisons test (with pooled variance) as post-hoc test and if not, a Brown-Forsythe ANOVA with Dunnett’s T3 multiple comparisons test (with individual variance calculated for each comparison) was used. If data were not normally distributed, lognormality was tested and in case the data were log-normally distributed, the aforementioned parametric statistical tests were performed on the log-transformed data. If data were not log-normally distributed, a non-parametric Kruskal-Wallis test was used on non-transformed data, with Dunn’s multiple comparisons test as post-hoc test. Differences between groups were considered statistically different when *P* < 0.05.

To assess the interrelationships between the expression levels of each gene and the developmental stage, we conducted a Principal Component Analysis (PCA) on standardized data and with principal components selected based on parallel analysis (confidence level of 95% and 1000 simulations). The contribution of each principal component to the total variance explained was calculated and plotted separately and cumulatively. From the selected principal components, the loadings and PC scores were plotted as well as a biplot showing the relation between these two plots. For the selected PCs, the correlation with each gene was calculated.

When comparing the effect of multiple factors (developmental stage & pH, pH & crispant status or crispant status & HEA/HEPES treatment) on nitrogenous waste excretion and gene expression levels, a two-way ANOVA was performed, with a full model fitted (including interaction effects). In cases where one or more factors had a significant effect (*P* < 0.05), a Tukey post-hoc test was used for multiple comparison testing.

## Results

### Characterization of larval nitrogen excretion and expression patterns of genes involved in nitrogen metabolism

We tracked zebrafish ammonia excretion patterns from embryo to the late larval stage and subsequently characterized the expression patterns of genes involved in nitrogen metabolism and ionocyte marker genes. Nitrogenous waste excretion was relatively low at 2-3 dpf (below 1 nmol per fish) and increased to approximately 4 nmol per fish at 5 dpf, with a decrease to approximately 2 nmol per fish afterwards (Fig. 1A). After the larval stage, *i.e.*, from around 20 dpf onwards, the total nitrogenous excretion per fish strongly increased to a maximum between 1 and 2 µmol per fish (Fig. S1A). We quantified both ammonia and urea excretion, and the contribution of these two nitrogenous waste products to total nitrogenous waste excretion shifted during larval development from urea to ammonia excretion (Fig. 1B), although a small percentage of nitrogen remains excreted as urea even beyond the larval stage (Fig. S1B).

**Fig. 1:**
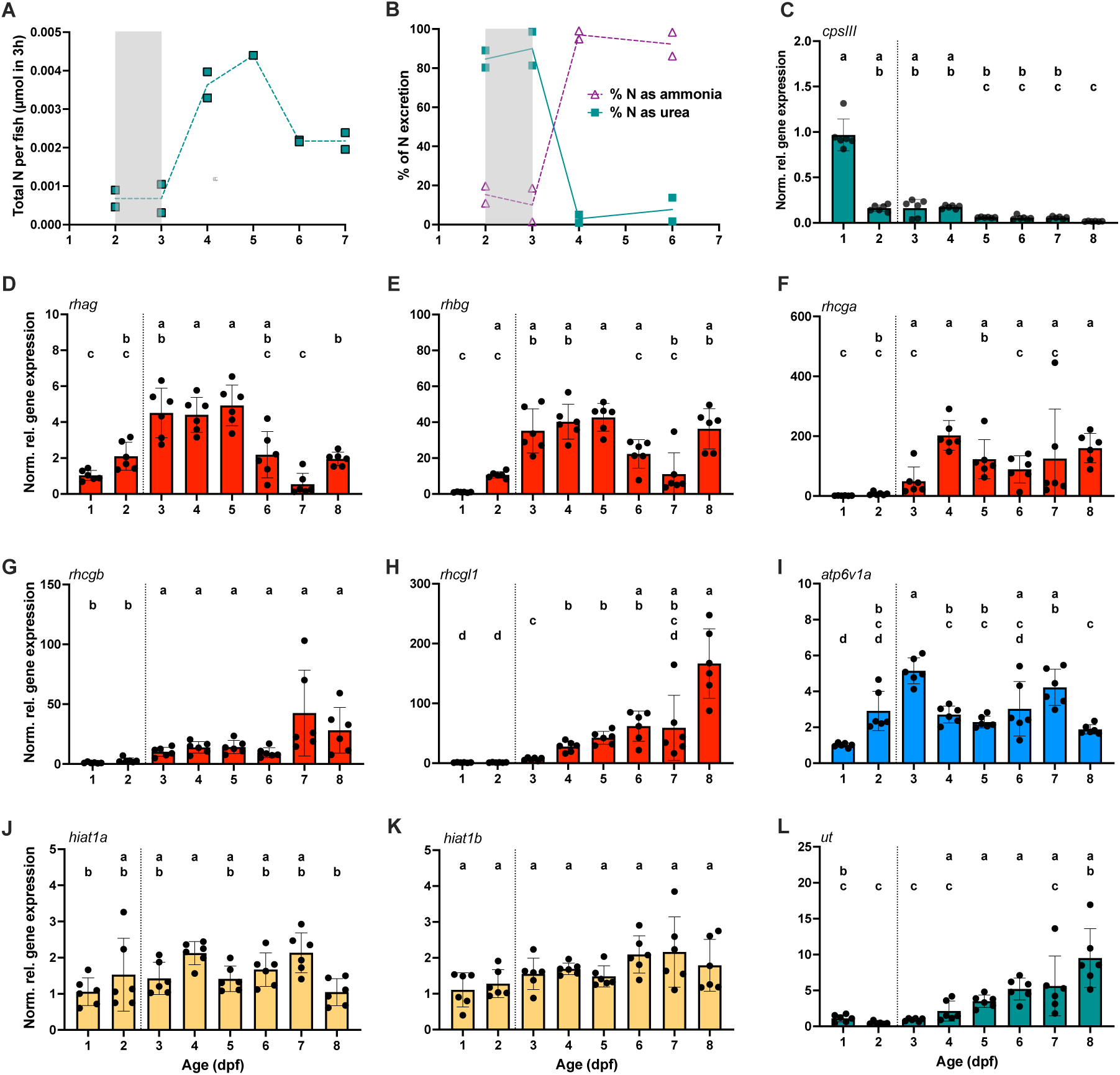
Characterization of zebrafish nitrogen excretion and expression patterns of genes involved in nitrogen metabolism from 1 to 8 days post-fertilization. **A**: Total nitrogen (in µmol N per fish in 3 hours) excreted by zebrafish larvae from 2-7 days post-fertilization (dpf). N-excretion was measured in duplicate for every dpf by measuring water ammonia and urea accumulation during 3 hours, with urea concentrations multiplied by 2 to obtain the corresponding µmol nitrogen. The time of hatching is indicated with a grey box between 2 and 3 dpf. B: Relative excretion of ammonia (open triangles) and urea (closed squares) in 2-7 dpf zebrafish. The time of hatching is indicated with a grey box between 2 and 3 dpf. C-K: Gene expression levels of nitrogen metabolic genes and ionocyte markers throughout zebrafish development (1-8 dpf). C: Normalized relative expression of carbamoyl-phosphate synthase iii. D-H: Normalized relative expression of rhesus glycoproteins (rhag, rhbg, rhcga, rhcgb and rhcgl1). I: Normalized relative expression of ionocyte marker H^+^-ATPase (atp6v1a) J-K: Normalized relative expression of alternative ammonia transporters hippocampal-abundant transcript 1a and 1b (hiat1a, hiat1b) L: Normalized relative expression of urea transporter ut. Gene expression was measured in pooled embryonal and larval samples (n = 6 per group). Gene expression is expressed as normalized relative gene expression (normalized to 1 dpf samples, and calculated compared to the relative expression of two housekeeping genes: elongation factor 1a (elf1a) and 40S ribosomal protein 11 (rps11). Individual values are shown for each group, with a colored bar and error bars indicating mean relative expression ±SD. Bar colors correspond to gene categories (teal: urea metabolism, red: rhesus glycoproteins, cyan: ionocyte-marker, yellow: hiat-ammonia transporters). Moment of hatching is indicated with a dotted line between 2 and 3 dpf. Significance between developmental stages was tested with a one-way ANOVA or non-parametric equivalent (depending on normal distribution of data) and appropriate post-hoc tests (Dunnett’s T3 test for parametric ANOVA and Dunn’s test for Kruskal-Wallis). Significant differences (p < 0.05) in relative gene expression are indicated using different lower-case letters.

Afterwards, we examined the whole-body expression of genes involved in nitrogen metabolism and ionocyte marker genes. Carbamoyl phosphate synthase III (*cpsiii*) expression was highest at 1 dpf and decreased significantly from 4 dpf onwards (Fig. 1C), which closely corresponds to the decrease in urea excretion. Other ornithine-urea cycle genes varied throughout development (Fig. S2). From hatching onwards (between 2 and 3 dpf), increases in transcript abundance of *rhag, rhbg, rhcga* and later *rhcgb* and *rhcgl1* were observed (Fig. 1D-H). Broadly, two expression patterns are apparent for rhesus glycoproteins: *rhag, rhbg* and *rhcga* increase 4-400-fold from hatch onwards while *rhcgb* and *rhcgl1* increase 50 to 200-fold at later developmental times (Figs. 1, S3). This suggests that under baseline conditions, the expression of *rhag*, *rhbg* and *rhcga* are involved in larval nitrogenous waste excretion, while *rhcgb* and *rhcgl1* expression increases at a later stage in development. Together with the increase in rhesus glycoprotein expression, an increase of the HR-ionocyte-marker H^+^-ATPase (*atp6v1a*) transcript abundance was observed (Fig. 1I). In contrast to the rhesus glycoproteins, *hiat1a* only increased 3-fold at 4dpf compared to 1 dpf, whereas *hiat1b* transcript abundance did not change throughout the developmental period (Fig. 1J-K). Finally, while urea excretion decreased over time, the expression of urea transporter (*ut*) increased in a similar pattern as *rhcgb* and *rhcgl1* (Fig. 1L).

Subsequently, we assessed whether the different whole-body expression patterns of rhesus glycoproteins correspond to a difference in localization. We performed whole-mount immunohistochemistry on 5 and 6 dpf larvae targeting Rhag, Rhbg and Rhcgb, together with a marker for HR-ionocytes (ConA) (Figs. 2 and 3). Consistent with the patterns of rhesus glycoprotein gene expression, we see a clear change in localization of HR-ionocytes from the yolk sac skin to the developing gill filaments at 5 dpf (Fig 2). Accordingly, rhesus glycoproteins also show developmental stage-specific localization. At 5 dpf, both the ConA signal and Rhag and Rhbg proteins are present on the gill filaments (Fig. 2A, B, D). While the yolk sac epithelium still has many ConA-positive cells, these cells are also present in the region of the developing gill filaments. In contrast, Rhcgb signal did not localize to the gill filaments, but was restricted to the yolk sac HR cells together with ConA signal (Fig. 2C, D). At 6 dpf, filament-expressed rhesus glycoproteins (Rhag, Rhbg) and the ionocyte-expressed Rhcgb are located in the gill area: Rhag and Rhbg show similar localization as at 5 dpf (Fig. 3A, B, D), while Rhcgb localizes to HR ionocytes between the developing filaments (Fig. 3C, D), with most cells showing signal from both ConA and Rhcgb antibody. From these data, it is evident that there are indeed two main pathways of rhesus-based ammonia excretion that appear in the gills at different moments. The first pathway, in early larval development (3-5dpf) involves Rhbg (and possibly Rhag) in developing branchial filaments. The second pathways occurs later in development when Rhcbg is expressed in gill ionocytes.

**Fig. 2:**
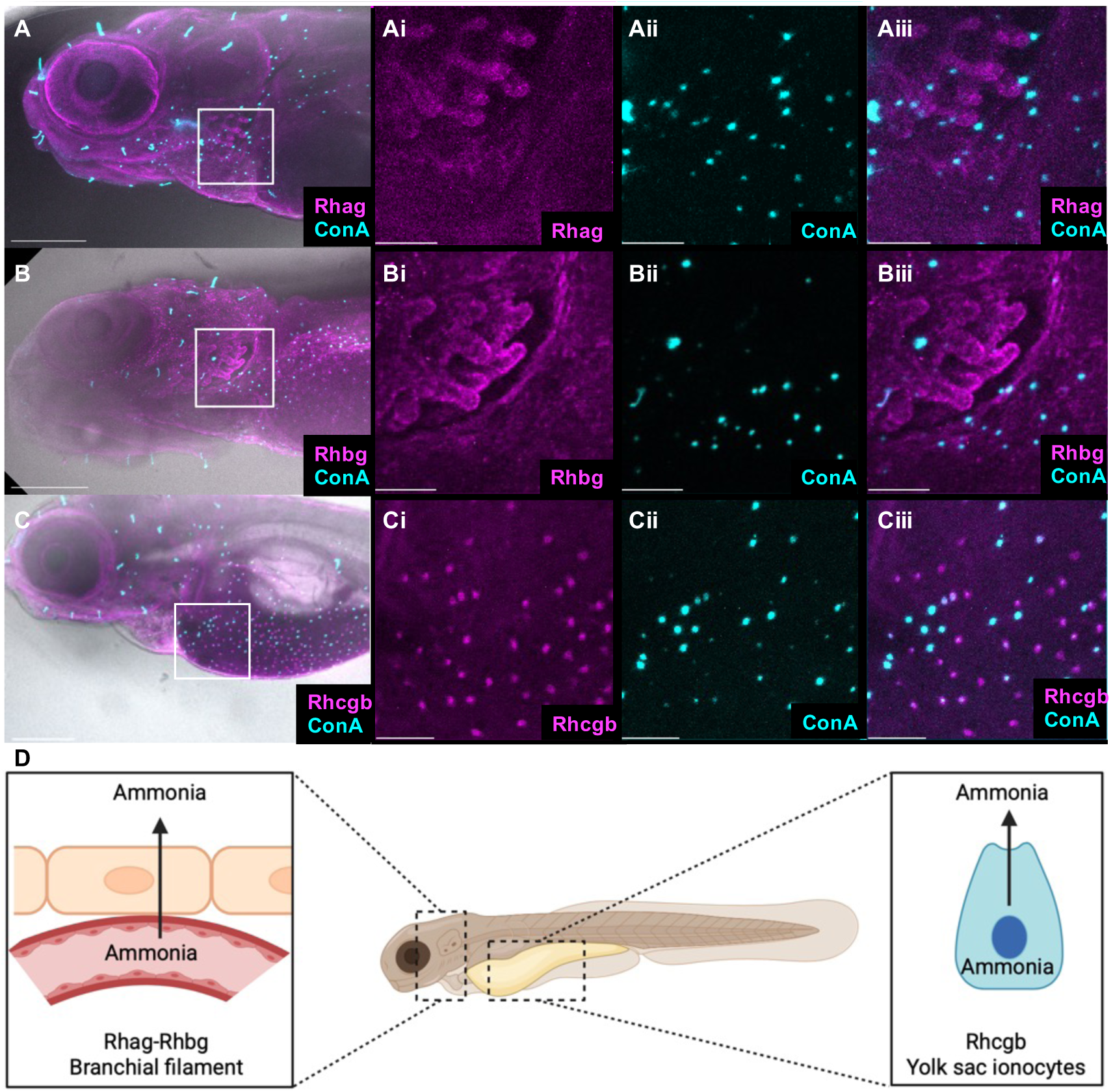
Immunofluorescence of rhesus glycoproteins and H^+^-ATPase-rich ionocytes in 5 dpf zebrafish. A) Rhag antibody (1:500) with concanavalin A to identify HR-ionocytes. A brightfield image is used for reference and fluorescent signals of Rhag-antibody (magenta) and concanavalin A (cyan) are both shown. Ai-Aiii inserts are detail images of the white square area: (Ai) Rhag signal on developing gill filaments, (Aii) ConA signal on developing gill filaments, (Aiii) Overlay of both channels. B) Rhbg antibody (1:250) with concanavalin A. A brightfield image is used for reference and fluorescent signals of Rhbg-antibody (magenta) and concanavalin A (cyan) are both shown. Bi-Biii inserts are detailed images of the white square area: (Bi) Rhbg signal on developing gill filaments, (Bii) ConA signal on developing gill filaments, (Biii) Overlay of both signals. C) Rhcgb antibody (1:500) with concanavalin A to identify HR-ionocytes. A brightfield image is used for reference and fluorescent signals of Rhcgb-antibody (magenta) and concanavalin A (cyan) are both shown. Ci-Ciii inserts are detail images of the white square area: (Ci) Rhcgb signal on the yolk sac epithelium, (Cii) ConA signal on yolk sac epithelium, (Ciii) Overlay of both channels showing partial overlap of fluorescent signal. For all overview images, the scale bar indicates 200 µm, for all inserts the bar indicates 50µm. D) Schematic representation of the localization of rhag, rhbg and rhcgb in different cell types in the yolk sac or gill of a larval zebrafish. Created in https://BioRender.com

**Fig. 3:**
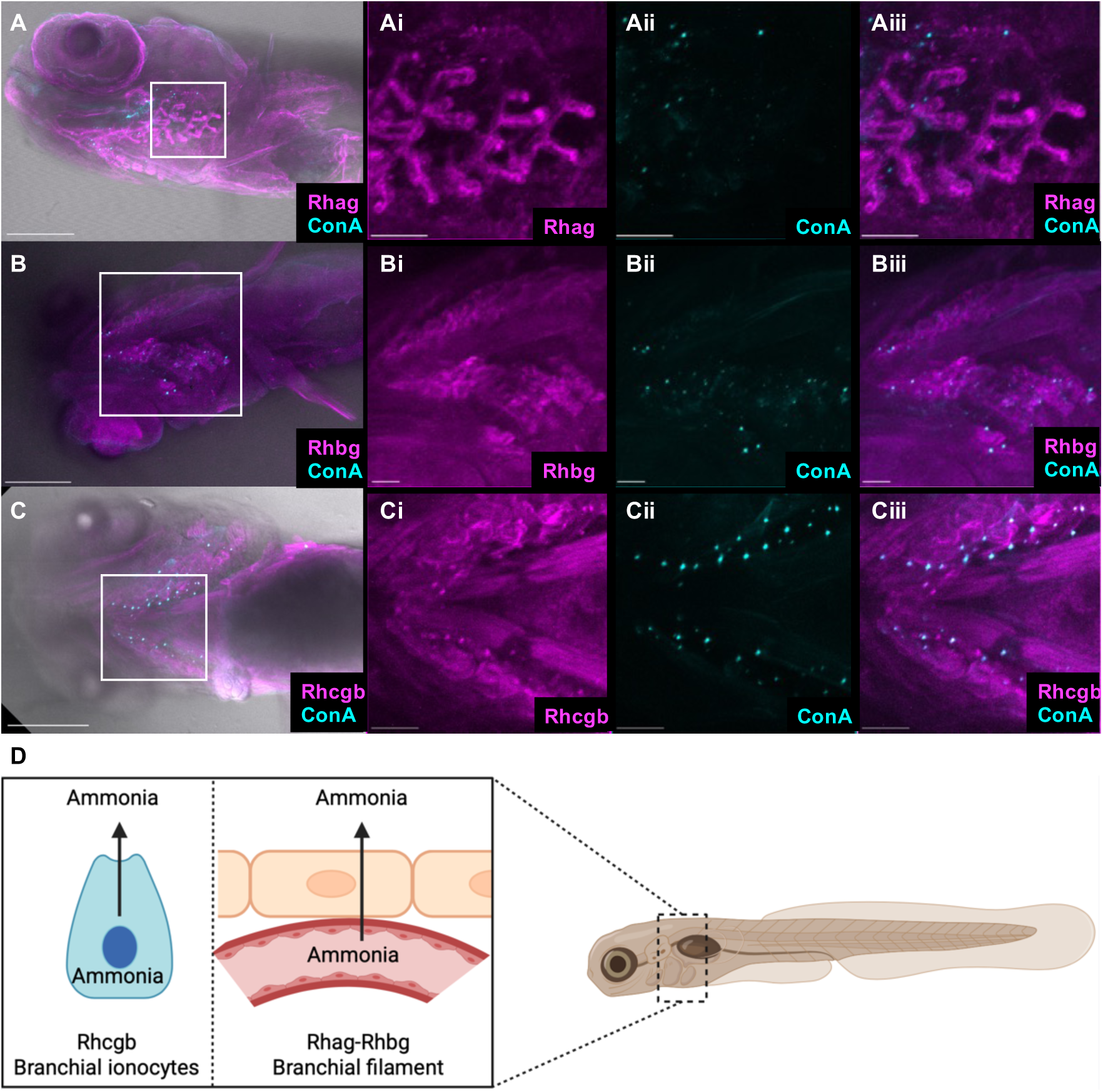
Immunofluorescence of rhesus glycoproteins and H^+^-ATPase-rich ionocytes in 6 dpf zebrafish. A) Rhag antibody (1:500) with concanavalin A to identify HR-ionocytes. A brightfield image is used for reference and fluorescent signals of Rhag-antibody (magenta) and concanavalin A (cyan) are both shown. Ai-Aiii inserts are detail images of the white square area: (Ai) Rhag signal on developing gill filaments, (Aii) ConA signal on developing gill filaments, (Aiii) Overlay of both channels. B) Rhbg antibody (1:250) with concanavalin A. A brightfield image is used for reference and fluorescent signals of Rhbg-antibody (magenta) and concanavalin A (cyan) are both shown. Bi-Biii inserts are detailed images of the white square area: (Bi) Rhbg signal on developing gill filaments, (Bii) ConA signal on developing gill filaments, (Biii) Overlay of both signals. C) Rhcgb antibody (1:500) with concanavalin A to identify HR-ionocytes. A brightfield image is used for reference and fluorescent signals of Rhcgb-antibody (magenta) and concanavalin A (cyan) are both shown. Ci-Ciii inserts are detail images of the white square area: (Ci) Rhcgb signal on developing gill filaments, (Cii) ConA signal on developing gill filaments, (Ciii) Overlay of both channels showing partial overlap of fluorescent signals. For all overview images, the scale bar indicates 200 µm, for all inserts the bar indicates 50µm. D) Schematic representation of the localization of rhag, rhbg and rhcgb in different cell types in the gill of a larval zebrafish. Created in https://BioRender.com

### Contribution of rhesus glycoproteins to nitrogen excretion at low environmental pH

Based on the two gene expression patterns and respective localizations of rhesus glycoproteins (cutaneous/epithelial *vs.* ionocyte localization), we hypothesized that these ammonia excretion pathways (epithelial and ionocyte-associated) provide larvae with the versatility to respond to challenging environmental conditions. To test this hypothesis, we examined changes in expression of rhesus glycoproteins in response to low pH, which was predicted to increase the ionocyte-expressed rhesus glycoproteins in particular due to its association with H^+^-ATPase and Na^+^/H^+^-exchanger 3b activity. The results of this experiment suggest that under low environmental pH, expression of of *rhag, rhbg* and *rhcga* at 5 dpf is increased 2-fold compared to levels in larvae raised at standard pH (7.4) (Fig. 4A-C). Transcript abundance of *rhcgb* and *rhcgl1* expression was unaffected (Fig.4D, E). We found no effect on the localization of Rhag, Rhbg or Rhcgb at 5 dpf, when larvae were reared at pH 5.0, compared to larvae raised at a pH of 7.4 (data not shown). This suggests that while localization remains unchanged, the larvae respond to low pH by upregulating expression of epithelial rhesus glycoproteins. *Atp6v1a* and *nhe3b* expression was elevated (Fig. 4F), in line with our hypothesis whereas *cpsIII* expression was decreased at both 4 and 5 dpf in response to a low pH (Fig. 4H) and *ut* expression was not affected by low pH (Fig. 4I), indicating that H^+^-ATPase and Na^+^/H^+^-exchanger 3b activity is likely higher to address the low pH effects on Na^+^-uptake, whereas urea production decreased.

**Fig. 4:**
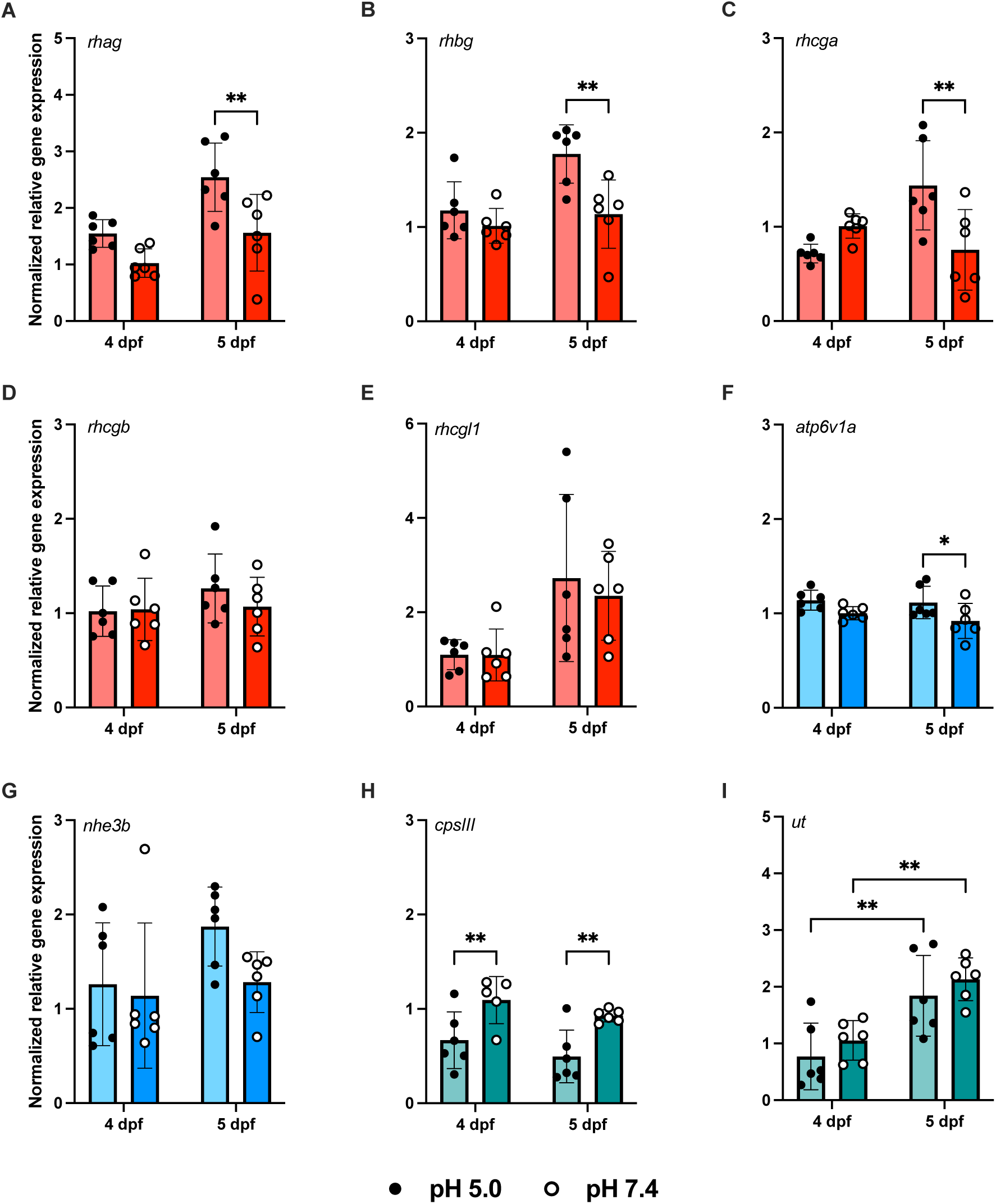
Effect of environmental pH on nitrogen metabolic and ionocyte-related gene expression at 4- and 5 days post-fertilization (dpf). Expression of rhesus glycoproteins (rhag, rhbg, rhcga, rhcgb and rhcgl1; A-E), ionocyte marker genes (H^+^-ATPase; atp6v1aa, Na^+^/H^+^-exchanger 3b; nhe3b F-G), and urea metabolism genes (cpsIII, ut (H-I) was measured in pooled embryonal and larval samples (n = 6 per group). Gene expression is expressed as normalized relative gene expression (normalized to 4 dpf, pH 7.4 samples, and calculated relative to the expression of two housekeeping genes: elongation factor 1a (elf1a) and 40S ribosomal protein 11 (rps11). Individual values are shown for each group, with a colored bar and error bars indicating mean relative expression ±SD. Bar colors correspond to gene categories (red: rhesus glycoproteins, cyan: ionocyte-markers, teal: urea metabolism), with the lighter shade indicating low pH and a darker shade indicating control pH. Zebrafish reared at pH 5.0 are shown with closed (black) circles, and those reared at pH 7.4 are shown with open (white) circles. Significance between developmental stages and environmental pH levels was tested with a two-way ANOVA and Tukey post-hoc tests. Asterisks indicate the significance level of Tukey post-hoc tests (* p < 0.05, ** p <0.01).

Having established differential responses of the epithelial- and ionocyte-linked rhesus glycoproteins to environmental pH, we next investigated how each pathway responds when the other is rendered non-functional. We measured nitrogen excretion as well as localization and expression of rhesus glycoproteins in F0-crispants for *rhbg* (epithelial) and *rhcgb* (ionocyte) at 5 dpf under neutral (7.4) and low (5.0) pH. Based on IHC, we confirmed an absence of both rhesus glycoproteins in their respective crispant groups (Fig. S4). This was, however, not associated with significant differences in nitrogen excretion, suggesting redundancy within the system, which has been reported before in *rhcgb*-KO zebrafish. Both *rhbg*- and *rhcgb*-crispants had similar total nitrogen excretion rates as controls (Fig. 5A) and while they produced less urea at pH 5.0, this response was also observed in controls (Fig. 5B).

**Fig. 5:**
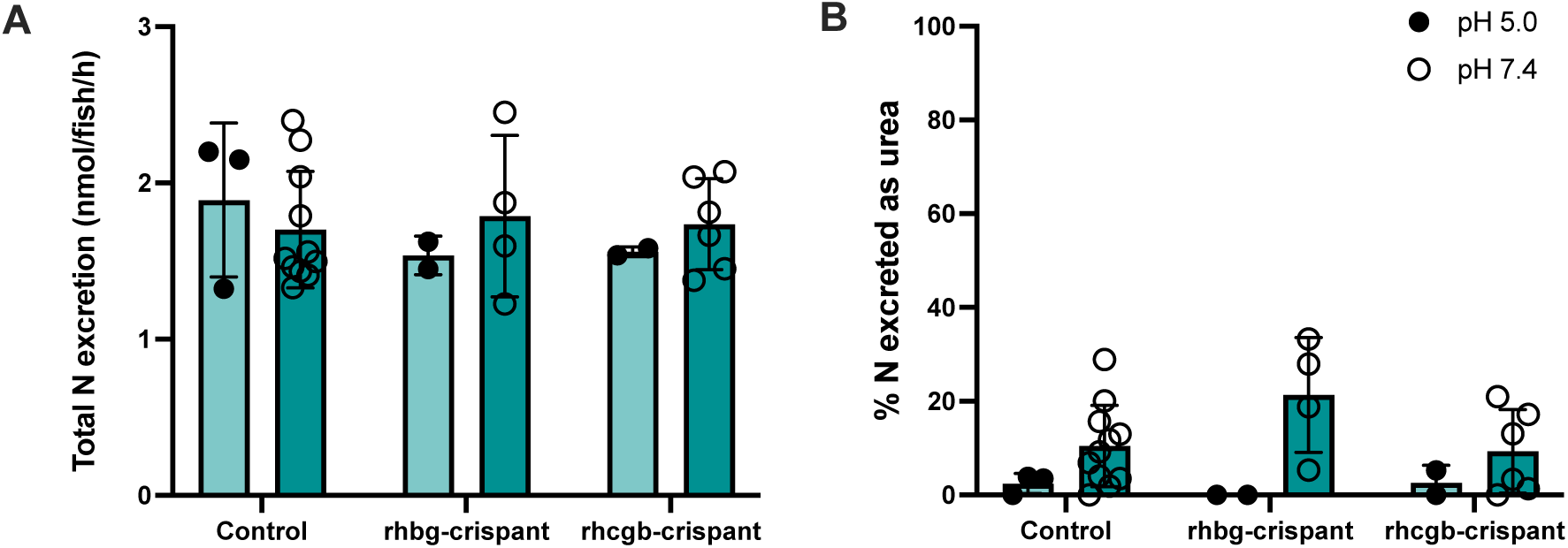
Nitrogenous waste excretion of control and crispant zebrafish larvae under different environmental pH levels. A: Total nitrogen (in nmol fish^-1^ h^-1^) excreted by zebrafish larvae. Total N excretion per fish was calculated by measuring water ammonia and urea accumulation during 3 hours, with urea concentrations multiplied by 2 to obtain the corresponding nmol nitrogen (urea contains 2 nitrogen molecules). **B:** Relative excretion of urea expressed as percentage of total N excretion. Each point was calculated from pooled larvae (10-11 fish per sample). Nitrogen waste excretion of zebrafish reared at pH 5.0 are shown with closed (black) circles, and those reared at pH 7.4 are shown with open (white) circles.

To determine whether there were compensation mechanisms between rhesus glycoproteins that would explain the lack of difference in ammonia excretion, we next looked at expression levels of these proteins in each crispant under low and normal pH (Fig. 6). *Rhbg*-crispants showed comparable responses to low pH as controls, with a significant increase in *rhcga* expression (Fig. 6C). However, they had lower expression of *rhbg* than the controls at both pH levels (Fig. 6B) and also showed no compensatory increase in other rhesus glycoprotein expression (*rhag*, *rhcgb* or *rhcgl1*) (Fig. 6A, D, E). In contrast, *rhcgb*-crispants did not exhibit an increase in rhesus glycoprotein expression at a low pH at all, and instead showed a significantly lower expression of all rhesus glycoproteins than control fish at pH 5.0, and a lower expression of *rhag*, *rhcgb* at pH 7.4 (Fig. 6A-E). In contrast, *Rhcgb*-crispants had significantly higher transcript abundances of another class of potential ammonia transporters: *hiat1a* expression was significantly higher than in control fish at pH 7.4 and *hiat1b* expression at both pH 5.0 and 7.4 (Fig. 6F & G). In line with the similarity of *rhbg-* and *rhcgb-*crispants and controls in terms of urea excretion patterns (Fig. 5B), *cpsiii* expression was not affected in either crispant group (Fig. 6H), and *ut* expression was reduced (Fig. 6I). Hence, under both pH 5.0 and 7.4, the loss of functionality of *rhbg* or *rhcgb* did not lead to changes in nitrogen metabolism nor to increases in other rhesus glycoprotein expression, suggesting that *hiat1a* or *hiat1b* may be capable of compensating for the loss of rhesus glycoproteins.

**Fig. 6:**
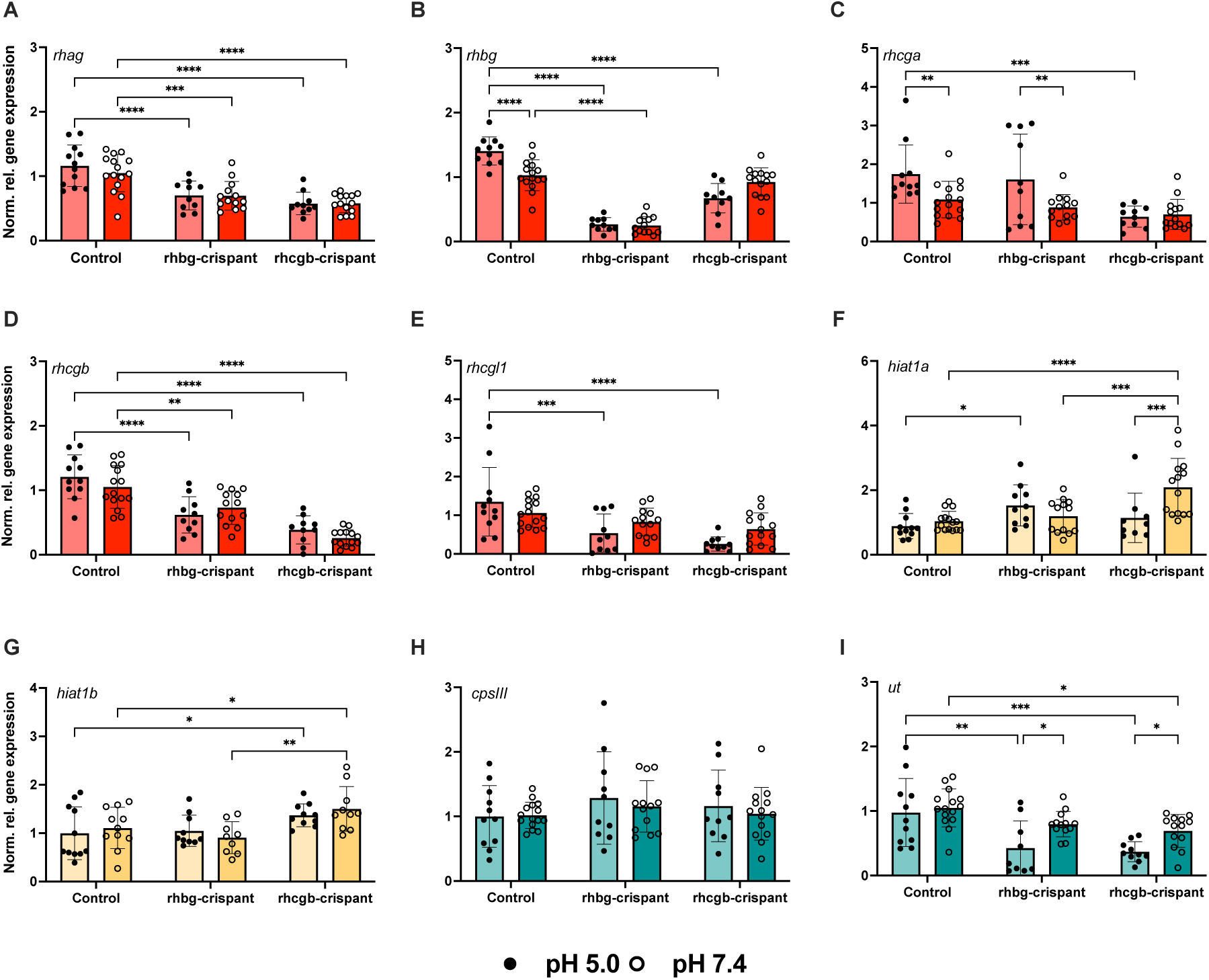
Effect of environmental pH on nitrogen metabolic and ionocyte-related gene expression in control, rhbg- and rhcgb-crispant zebrafish. Expression of rhesus glycoproteins (rhag, rhbg, rhcga, rhcgb and rhcgl1; A-E), hippocampal-abundant transcripts 1a and 1b (hiat1a, hiat1b; F-G) and urea metabolism genes (cpsIII, ut (H-I) was measured in pooled embryonal and larval samples (n = 9-15 per group). Gene expression is expressed as normalized relative gene expression (normalized to control fish reared at pH 7.4 and calculated relative to the expression of two housekeeping genes: elongation factor 1a (elf1a) and 40S ribosomal protein 11 (rps11). Individual values are shown for each group, with a colored bar and error bars indicating mean normalized relative expression ± SD. Colors correspond to gene categories (red: rhesus glycoproteins, yellow: hiat-ammonia transporters, teal: urea metabolism), with the lighter shade indicating low pH and a darker shade indicating control pH.. Zebrafish reared at pH 5.0 are shown with closed (black) circles, and those reared at pH 7.4 are shown with open (white) circles. Significance between different crispants and environmental pH levels was tested with a two-way ANOVA and Tukey post-hoc tests. Asterisks indicate the significance level of Tukey post-hoc tests (* p < 0.05, ** p <0.01), *** p < 0.001, **** p < 0.0001).

### Contribution of rhesus glycoproteins to nitrogen excretion during ammonia-related challenges

Since *rhcgb*-crispants showed a marked reduction in rhesus glycoproteins while retaining normal nitrogen excretion patterns, we further characterized the response of these animals to ammonia-related challenges (high external ammonia (HEA) and strong buffering of the medium with HEPES to inhibit the acid-trapping mechanism which enhances ammonia excretion). Exposure to HEA (500 µM NH_4_Cl) for three hours significantly decreased nitrogen excretion in both control larvae and *rhcgb-*crispants (Fig. 7A). However, while control larvae excreted their nitrogenous waste predominantly as ammonia, *rhcgb-*crispants excreted up to 80% of their nitrogenous waste as urea (Fig. 7B). In contrast, exposure to 10 mM HEPES-buffered embryo medium did not affect total nitrogenous waste excretion in control larvae and *rhgcb-*crispants (Fig. 7C), and increased urea excretion in both groups (Fig. 7D).

**Fig. 7:**
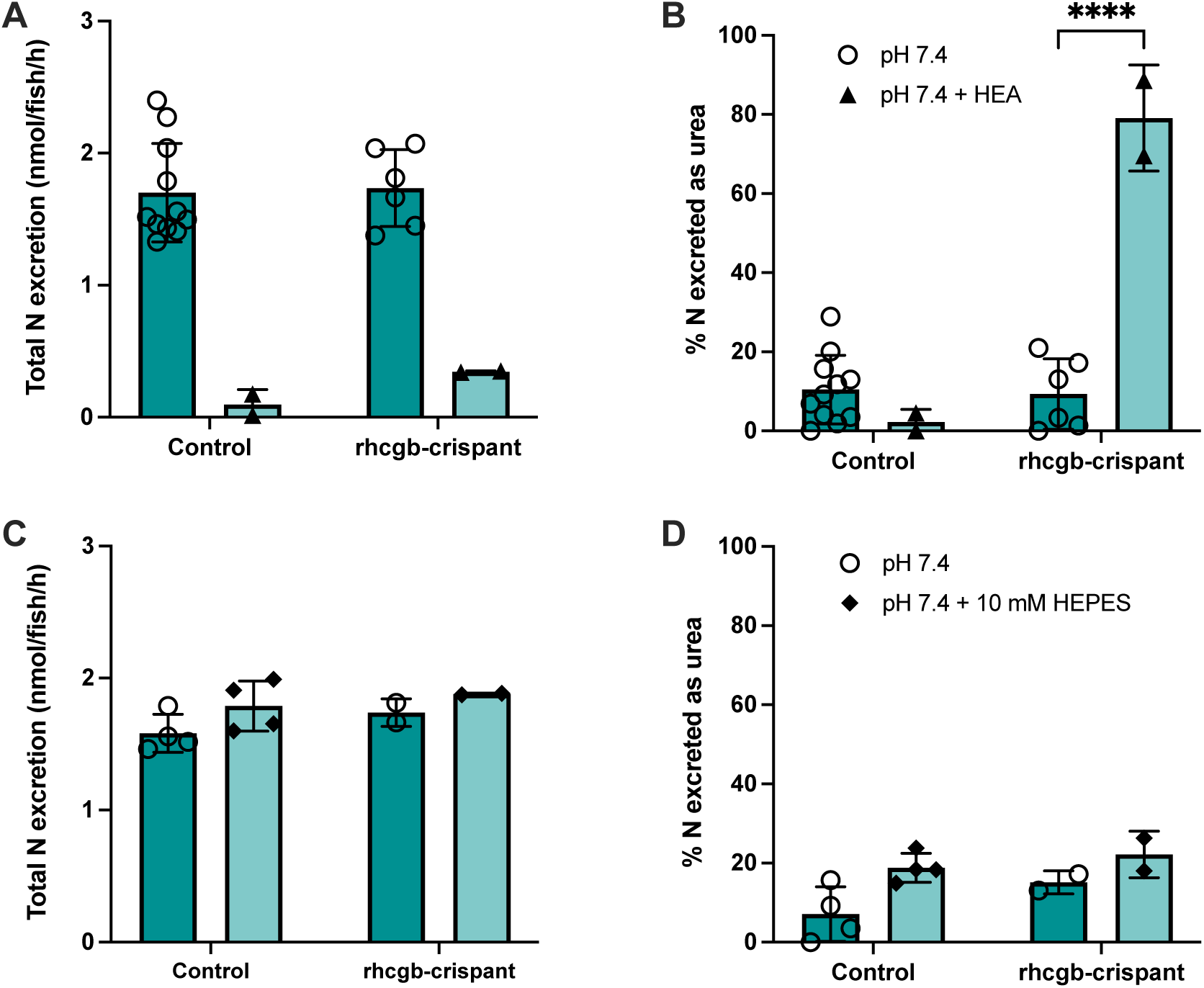
Nitrogenous waste excretion of control and rhcgb-crispant zebrafish larvae under ammonia-related challenges. A: Total nitrogen (in nmol fish^-1^ h^-1^) excreted by zebrafish larvae exposed to high external ammonia (500 µM NH4Cl) for 3 hours. Total N excretion per fish was calculated by measuring water ammonia and urea accumulation during 3 hours, with urea concentrations multiplied by 2 to obtain the corresponding nmol nitrogen (urea contains 2 nitrogen molecules). **B:** Relative excretion of urea expressed as percentage of total N excretion as calculated in A. Nitrogenous waste excretions of control zebrafish are shown as open circles, while those of rhcgb-crispants are shown as closed triangles. **C:** Total nitrogen (in nmol fish^-1^ h^-1^) excreted by zebrafish larvae exposed to 10 mM HEPES-buffered embryo medium for 3 hours. **D:** Relative excretion of urea expressed as percentage of total N excretion as calculated in C. Nitrogenous waste excretions of control zebrafish are shown as open circles, while those of rhcgb-crispants are shown as closed diamonds. Every point is calculated from pooled larvae (10 fish per sample). Significance between crispants and controls as well as the effect of exposure treatments were tested with a two-way ANOVA and Tukey post-hoc tests. Asterisks indicate the significance level of Tukey post-hoc tests (**** p < 0.0001).

Having established that *rhcgb*-crispants exhibited a switch in nitrogen excretion from ammonia to urea in response to HEA, we examined whether whole-body gene expression patterns explain this drastic change in nitrogenous waste excretion (Fig. 8). In control fish, exposure to 500 µM NH_4_Cl for 3 hours led to increased transcript abundances of genes involved in ammonia excretion: *rhag, rhbg, rhcga* and *rhcgb* were all significantly elevated (Fig. 8A-D), as were HR-ionocyte markers *atp6v1a* and *nhe3b* (Fig. 8F-G) and *hiat1b* (Fig. 8I). While *rhcgl1* and *hiat1a* were not elevated, these results indicate a strong transcriptional response to HEA exposure involving multiple pathways (epithelial and ionocyte-associated rhesus glycoproteins and *hiat*-mediated ammonia transport).

**Fig. 8:**
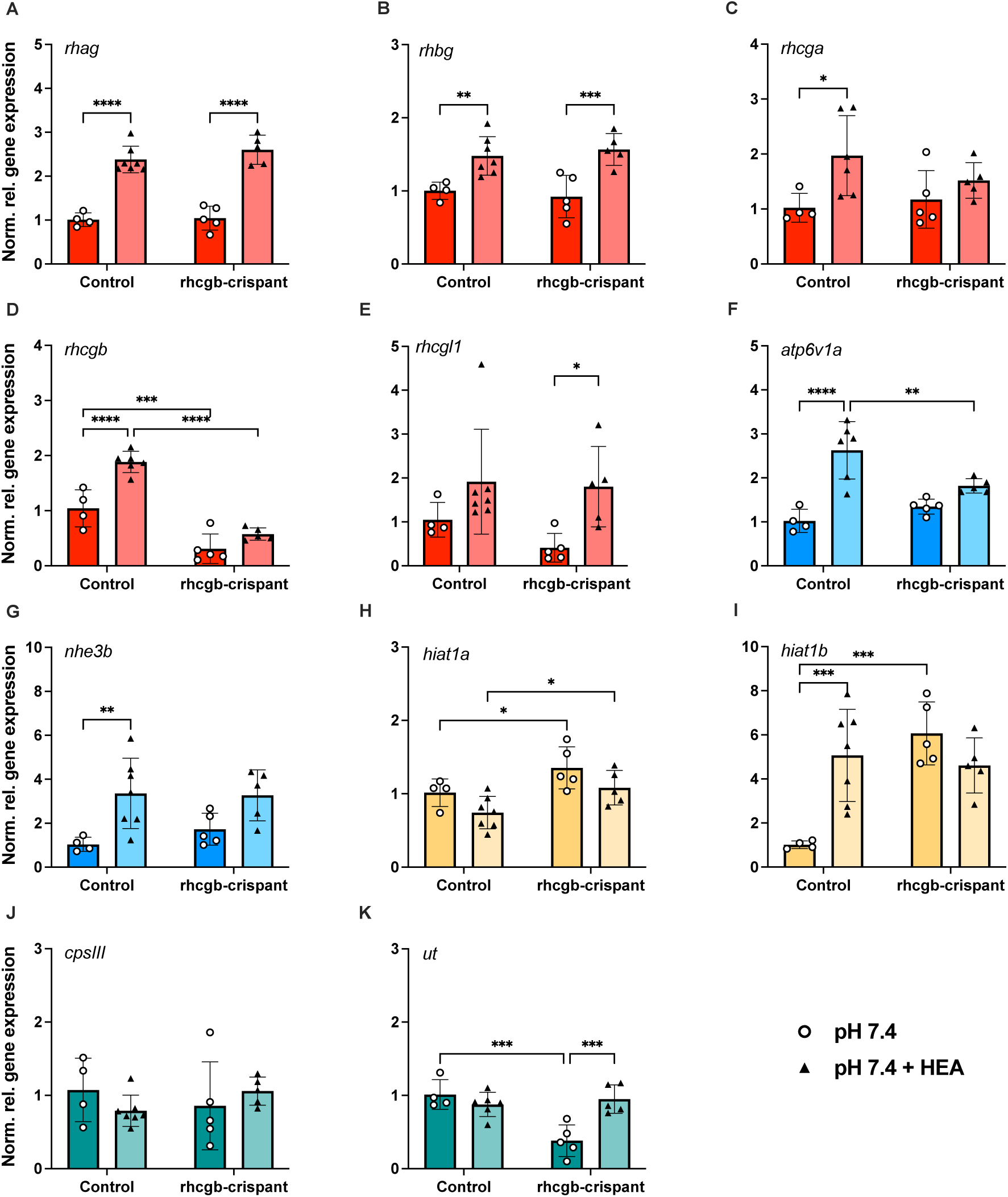
High external ammonia (HEA) alters nitrogen metabolic and ionocyte-related gene expression in 5 dpf control and rhcgb-crispant zebrafish. Control zebrafish and crispants were exposed to 500 µM NH4Cl for 3 hours and whole-body expression levels of nitrogen metabolic genes and ionocyte-markers were measured. Expression of rhesus glycoproteins (rhag, rhbg, rhcga, rhcgb and rhcgl1; A-E), HR-ionocyte markers (atp6v1aa, nhe3b; F-G), hippocampal-abundant transcripts 1a and 1b (hiat1a, hiat1b; H-I) and urea metabolism genes (cpsIII, ut (J-K) was measured in pooled embryonal and larval samples (n = 5-7 per group). Gene expression is expressed as normalized relative gene expression (normalized to unexposed control fish samples and calculated relative to the expression of two housekeeping genes: elongation factor 1a (elf1a) and 40S ribosomal protein 11 (rps11). Individual values are shown for each group, with a colored bar and error bars indicating mean relative expression ± SD. Bar colors correspond to gene categories (red: rhesus glycoproteins, cyan: ionocyte-marker, yellow: hiat-ammonia transporters, teal: urea metabolism). Unexposed zebrafish reared at pH 7.4 are shown as open circles, while zebrafish exposed to HEA are shown as closed triangles. Significance between crispants and control fish and between HEA-exposed fish and control conditions was tested with a two-way ANOVA and Tukey post-hoc tests. Asterisks indicate the significance level of Tukey post-hoc tests (* p < 0.05, ** p <0.01), *** p < 0.001, **** p < 0.0001).

In contrast, *rhcgb-*crispants responded to HEA only by upregulating transcript abundance of three rhesus glycoprotein genes: *rhag, rhbg*, and *rhcgl1* (Fig. 8A, B, E). In addition, the crispants had a 6-fold higher *hiat1b* expression under baseline conditions, which was not further increased in response to HEA (Fig. 8I). Transcript levels of urea transporter *ut* were also significantly elevated in *rhcgb*-crispants (Fig. 8K), in line with the higher percentage of nitrogen excreted as urea (Fig. 7B).

Three-hour exposure to 10 mM HEPES resulted in similar rhesus glycoprotein expression patterns in control larvae, with significant increases in *rhag*, *rhcga, rhcgb*, and *rhcgl1*, as well as ionocyte-markers (Fig. S5A-G) but did not affect *hiat1a* or *hiat1b* expression (Fig. S5J-K). *Cpsiii* and *ut* expression was also increased (Fig. S5H-I), in line with the significant increase in urea excretion in HEPES-exposed control fish (Fig. 7D). *Rhcgb-*crispants did not exhibit increased *rhag* and *rhcgb* expression in response to 10 mM HEPES exposure, but otherwise showed generally similar trends in rhesus glycoprotein expression, with elevated expression of *rhcga* and *rhcgl1* compared to *rhcgb-*crispants reared in control conditions (Fig. S5C, E). However, the change in expression of *rhcga* was lower in crispants than in control animals, and *rhcgb*-crispants also showed no increases in *atp6v1a* and *ut* expression. In contrast to what was seen in the HEA exposure experiment, *hiat1a* and *hiat1b* expression was not elevated in crispant zebrafish. HEPES-exposure thus did not affect nitrogenous waste excretion to the same extent as HEA-exposure and generally *rhcgb*-crispant larvae were able to retain similar ammonia and urea excretion patterns as control larvae.

## Discussion

In the present study, we have expanded current knowledge of zebrafish larval nitrogen metabolism by studying two distinct rhesus glycoprotein-mediated ammonia excretion pathways (summarized by Zimmer, 2024) and show that they exhibit different dynamics during the developmental skin-to-gill transition. Furthermore, we show that loss of either pathway alone does not impair ammonia excretion under baseline conditions or at low pH, and that neither pathway directly compensates for the loss of the other. Instead, *rhcgb* crispants upregulate expression of the alternative ammonia transporter *hiat1b*. Despite an apparently normal nitrogen excretion profile, *rhcgb* crispants exhibit impaired ammonia excretion under high external ammonia and shift toward urea excretion. Together, our findings indicate that zebrafish larvae possess a versatile and robust nitrogen excretion system at 5 dpf that maintains overall homeostasis even when individual components are disrupted.

In the current study, we found that the two spatially separate pathways of rhesus glycoprotein-mediated ammonia excretion (keratinocyte- and ionocyte-localized) also differ temporally in their skin-to-gill transition. The expression of Rhbg (as well as Rhag) in gill cells was apparent from 5 dpf onwards. This localization in the gill is similar to that found by Braun et al. (2009a) at 8 dpf using *in situ* hybridization, and in more detailed analyses of adult zebrafish gills (Porteus et al., 2021), Rhbg was found on the gill epithelial cells. The expression of Rhcgb was clearly distinct and colocalized with the HR-ionocyte marker ConA on both the yolk sac (until 5 dpf) and between the gill filaments (from 6 dpf onwards). Previously, its localization in the gill has been confirmed in adults with immunohistochemistry and in 8 dpf larvae with *in situ* hybridization before (Nakada et al., 2007a; Braun et al., 2009a).

To our knowledge, this is the first time that a difference in timing of the skin-to-gill transition of rhesus glycoproteins has been reported. Together with the rapid increase in ammonia excretion from hatching onwards, these findings show that the gill becomes necessary for ammonia excretion early in zebrafish development, similar to what has been reported for rainbow trout (*Oncorhynchus mykiss*) larvae (Zimmer et al., 2014; Zimmer and Wood, 2015). This is in line with the ionoregulatory hypothesis for gill development, where the gill is needed for osmoregulation, particularly Na^+^-uptake, before it plays a role in oxygen uptake (Rombough, 2002; 2007). Ammonia excretion enables Na^+^-uptake via the metabolon of Rhcgb, H^+^-ATPase and Na^+^/H^+^-exchanger 3b (Wright and Wood, 2009). This would explain the localization of Rhcgb in the gill from 6 dpf onwards. For rainbow trout, it was shown that the skin-to-gill transition of ammonia excretion occurs regardless of its role in Na^+^-uptake (Zimmer and Wood, 2015). The earlier appearance of Rhbg expression in the gill of zebrafish would also support this ‘ammonia-first’ hypothesis (Zimmer et al., 2017), since its function is independent of ionoregulation as far as we currently know.

In line with the correlation between ammonia excretion rates and rhesus glycoprotein expression, we found that a low environmental pH leads to increased ammonia (compared to urea) excretion. In addition, it resulted in upregulated transcription of rhesus glycoprotein genes (*rhag* and *rhbg*) that are expressed in the gills by this moment according to our results. Expression of *rhcgb* was unaffected by rearing at pH 5.0, which leads to the question whether its lack of response is due to the reduced need of ionocyte-mediated ammonia excretion, which requires active H^+^-export via H^+^-ATPase, or due to the localization in the skin rather than gills. This question may be answered by extending the low pH exposure to 6-7 dpf, in line with the later appearance of *rhcgb* expression in the gills.

The pH-dependent increase in *rhbg* expression (localized to epithelial cells), but not the HR-ionocyte-linked rhesus glycoprotein expression shows that there are multiple pathways to excrete ammonia, and that they respond differently to environmental conditions. What is also clear is that zebrafish larvae show a high flexibility in their nitrogenous waste excretion: urea excretion was significantly reduced in fish reared at pH 5.0 and expression of the OUC marker *cpsiii* was significantly reduced at both 4 and 5 dpf. This versatility of zebrafish larvae in response to challenges to their physiological ammonia excretion mechanisms was also apparent in the crispants of *rhbg* and *rhcgb* we produced and characterized here. Despite lacking one pathway of ammonia excretion, total nitrogenous waste excretion was no different from control fish under normal or low pH, high external ammonia, or HEPES-buffered medium exposure. This underlines the fundamental and existential importance of dealing with nitrogenous waste and the evolved robustness of this system to external disturbances.

While both *rhbg*- and *rhcgb*-crispants were unaffected in nitrogen excretion compared to controls, our characterization revealed intriguing differences between the zebrafish that lacked *rhbg* compared to the ones that lacked *rhcgb* functionality. *Rhbg*-crispants showed similar expression patterns in general as controls at pH 7.4, and limited upregulation of *rhcga* transcript levels at pH 5.0. In contrast, *rhcgb*-crispants had significantly lower expression levels of all rhesus glycoproteins at both rearing pH levels. This finding is seemingly counterintuitive and suggests that loss of *rhcgb* affects the other rhesus glycoproteins negatively, instead of a compensatory upregulation of other rhesus glycoproteins, as shown by Zimmer and Perry (2020) for *rhcgb*-knockout zebrafish. Interestingly, our *rhcgb*-crispants showed an increased expression of the *hiat1a* and *hiat1b* ammonia transporters instead, in particular at neutral pH. Knockdown of *hiat1b* decreased ammonia excretion particularly at the yolk sac and pharyngeal region of 4 dpf zebrafish (Zhouyao et al., 2022) and our results support its role as an additional ammonia transporter that may take over the role of rhesus glycoproteins in *rhcgb* crispant zebrafish.

In addition, we compared the responses of the *rhcgb* crispants to short-term HEA- exposure and strong HEPES-buffering of the embryo medium with those of control larvae. HEA-exposure induced upregulation of both non-ionocyte-associated rhesus glycoproteins (*rhag, rhbg, rhcga*) and HR-ionocyte-associated glycoprotein (*rhcgb*) expression in control animals, similarly to findings from previous work (Braun et al., 2009b). Genes encoding other HR-ionocyte markers (H^+^-ATPase, Na^+^/H^+^-exchanger 3b) also responded to HEA-exposure, suggesting a concerted HEA-response in HR-ionocytes. Moreover, *hiat1b* expression increased 6-fold in response to HEA-exposure, in line with its proposed role as ammonia transporter (Zhouyao et al., 2022). This is the first time its response to an ammonia-related stressor has been shown in zebrafish, although as stated above, its recent discovery and lack of exact cellular localization makes the underlying mechanism in which it may contribute difficult to determine.

The *rhcgb*-crispants, by contrast, responded with increased *rhag, rhbg* and *rhcga* expression but lacked a response in *rhcgb* and other HR-ionocyte markers, suggesting that these cells do not respond to HEA when *rhcgb* is lost. Expression of *hiat1b* also did not increase further, which suggests a limit to the amount of ammonia excretion compensation through this pathway. In line with this hypothesis, urea excretion almost entirely replaced ammonia excretion in these animals during HEA-exposure, together with an increase in urea transporter expression. This suggests the induction of detoxification mechanisms and an inability to upregulate ammonia excretion mechanisms further. It also shows that the OUC is not inactivated by 5 dpf and that HEA can indeed induce increased OUC activity (Zimmer et al., 2017). These results again illustrate the highly versatile nitrogenous waste excretion and detoxification mechanisms in larval zebrafish.

In contrast to HEA-exposure, HEPES-exposure, which removes the acidified boundary layer and inhibits ammonia excretion (Wright et al., 1989; Shih et al., 2008), did not affect *rhcgb*-crispants differently than control animals. Both groups increased urea excretion, but ammonia excretion in HEPES-treated animals was not significantly altered, despite a lack of upregulation of *rhag, rhcgb, hiat1a and hiat1b,* and an attenuated increase in *rhcga* expression. Both control and *rhcgb*-crispant larvae had higher transcript abundance of *rhcgl1*, which has not been directly implicated in ammonia transport yet. However, older studies targeting *rhcgb* with morpholino knockdown are now known to have unintentionally targeted *rhcgl1* instead (Zimmer and Perry, 2020). Interestingly, these studies showed reduced ammonia excretion as a result of this *rhcgl1* knockdown (Braun et al., 2009a; Kumai and Perry, 2011), suggesting that it plays a role in this response, and that *rhcgb-*crispant zebrafish may maintain ammonia excretion through *rhcgl1* upregulation.

## Conclusion

Taken together, our study gives further support to the current ammonia excretion model of larval zebrafish skin that involves (at least) two distinct rhesus glycoprotein-mediated pathways that are spatially separate. We extend this by showing that both pathways respond differently to environmental conditions at 5 dpf, and that expression of the rhesus glycoproteins involved in both pathways moves to the gill at different developmental stages. Using crispant zebrafish targeting either pathway, we additionally show that baseline nitrogen excretions are similar to control zebrafish, but that the two pathways do not directly compensate for each other. Instead, *rhcgb*-crispants compensate via upregulating the expression of HIAT ammonia transporters, and when exposed to high external ammonia concentrations, they switch to urea excretion. Overall, our results show the versatility of the zebrafish larval nitrogen excretion system and its resilience to loss of parts of the system.

## Acknowledgments

The authors want to thank Antoon van der Horst, Kimberley Janssen and Jeroen Boerrigter for excellent zebrafish care. We thank Jan Zethof for help with gene expression measurements and Jelle Postma for his help with confocal microscopy and analysis of microscopic images.

## Data availability

Data and resource availability: All relevant data and details of resources can be found within the article and its supplementary information.

