## Supplementary data for "Zebrafish larval nitrogen excretion is flexible and resilient to loss of rhesus glycoproteins"

### Supplementary files

***Table S1: Primer sequences used for genotyping F0-crispant zebrafish.***

| Exon | Forward primer (5'-3') | Reverse primer (5'-3') | Amplicon size (bp) |
|---|---|---|---|
| *Rhbg* 1 | GATGATACCGATGCCAAGAAGT | TTCATCAGCAGTTCAACCAATC | 256 |
| *Rhbg* 2 | ATTTAACTTTCTCATCGCAGCC | TGTGTGCTTTCTTATGGGAACA | 281 |
| *Rhbg* 4 | TGAGCTGCAGTATGTAAAAGGTG | AAGAGGTCAGAGTGGTAGACGG | 239 |
| *Rhcgb* 1 | CCAACATCAGACTCAGTTTACCC | GAAAACATGCAAGGAATATTGTTCT | 286 |
| *Rhcgb* 2 | GGTTTTCAGGATGTTCATGTCA | CCGATCTTGATCTTTCCATCA | 182 |
| *Rhcgb* 3 | TTCATGCATGTGAAACATGGTA | GATCAGACTCACGTTGAGGACA | 277 |

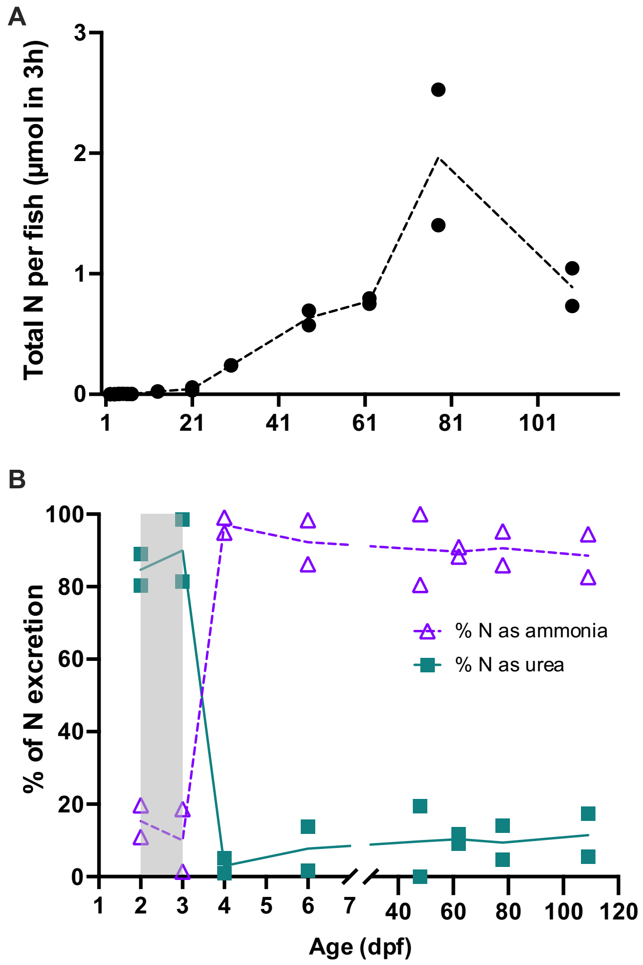


***Fig. S1: Nitrogen excretion of zebrafish across development. A:*** *Total nitrogen (in µmol N) excreted by zebrafish from all developmental stages (2-109 dpf). Total N excretion per fish was measured in duplicate for every day post-fertilization by measuring water ammonia and urea accumulation during 3 hours, with urea concentrations multiplied by 2 to obtain the corresponding µmol nitrogen (urea contains 2 nitrogen molecules). Duplicate values are plotted and a connecting line is plotted between mean values of the two duplicates.* ***B:*** *Relative excretions of ammonia (open triangles) and urea (closed squares) by zebrafish across development (2-109 dpf), expressed as percentage of total N excretion. Duplicate values are plotted per dpf, with a connecting line between mean values of the two duplicates.*

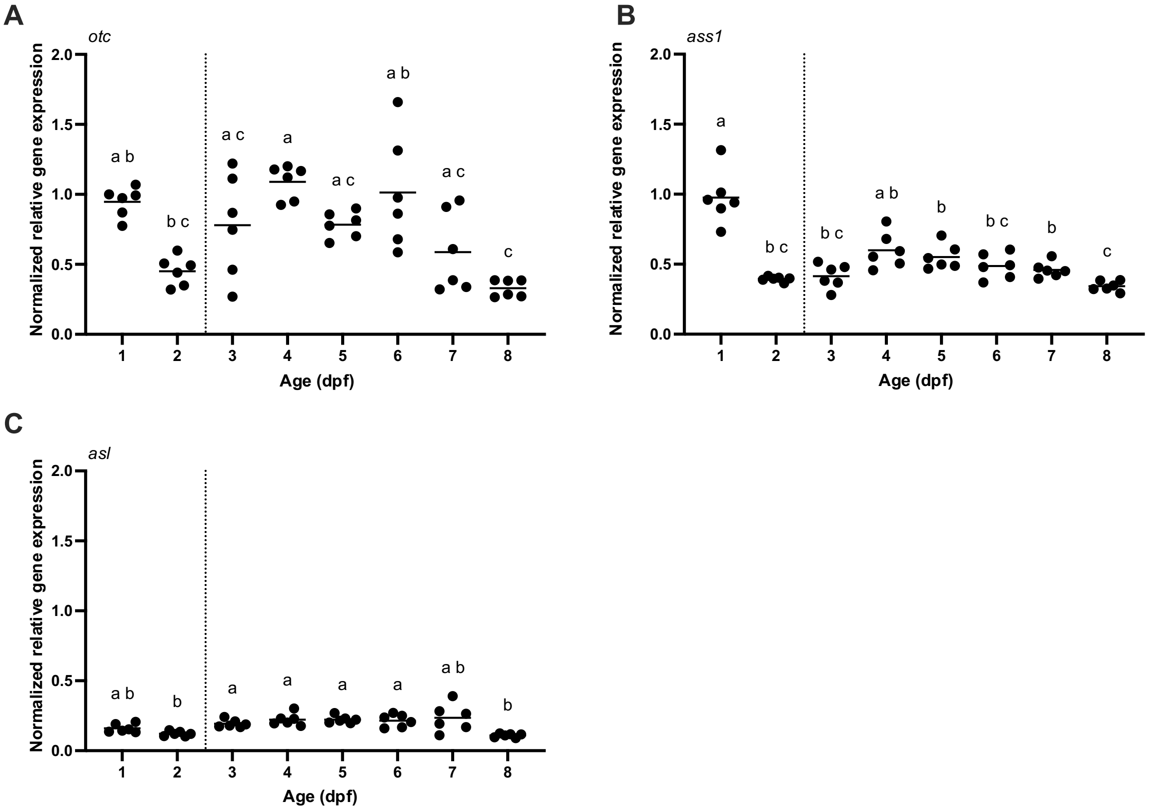


***Fig. S2: Expression of ornithine-urea cycle (OUC) genes in developing zebrafish larvae from 1 to 8 days post-fertilization.*** *Relative expression of ornithine transcarbamylase; otc (**A**), argininosuccinate synthetase 1; ass1 (**B**) and argininosuccinate lyase; asl (**C**) was measured in pooled embryonal and larval samples (n = 6 per group). Gene expression is expressed as normalized relative gene expression (normalized to the expression two housekeeping genes: $\beta$-actin and 40S ribosomal protein 11 (RPS11). Individual values are shown for each group, with a bar indicating mean relative expression. To make comparison between genes easier, y-axes are all the same scale. Moment of hatching is indicated with a dotted line between 2 and 3 dpf. Significance between developmental stages was tested with a one-way ANOVA or non-parametric equivalent (depending on normal distribution of data) and appropriate post-hoc tests (Dunnett's T3 test for parametric ANOVA and Dunn's test for Kruskal-Wallis). Significant differences ($p < 0.05$) in relative gene expression are indicated using different lower-case letters.*

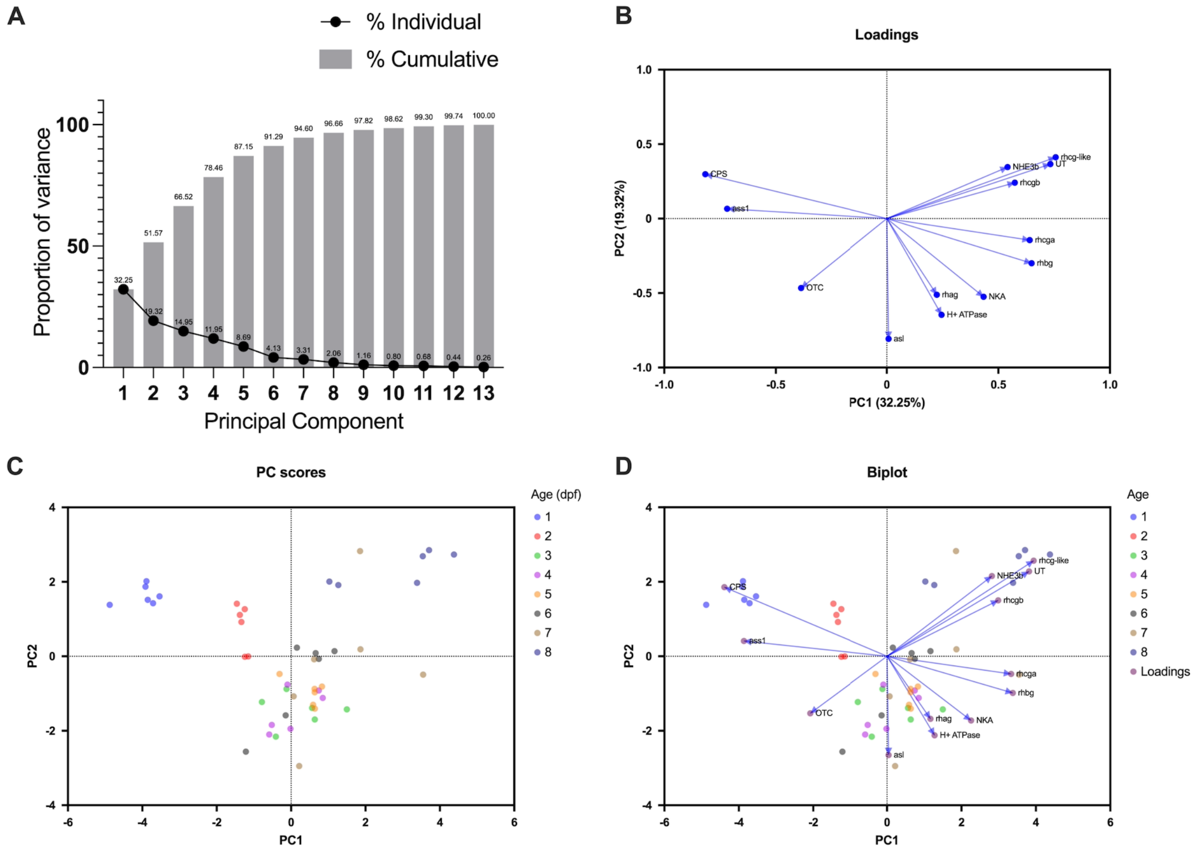


***Fig. S3: Principal component analysis of zebrafish nitrogen metabolism and excretion genes throughout development. (A):*** *Proportion of variance plot of PCA. The line shows the proportion of total variance explained for each PC. The grey bars indicate the cumulative variance explained by all PCs up to that point.* ***(B):*** *Loadings plot of PCA, showing correlations between expression patterns of each gene investigated.* ***(C):*** *Principal component scores plot of PCA, showing clustering of zebrafish samples (n= 48) on the first two principal components. Samples are colored by developmental age (in dpf).* ***(D)****: Biplot of PCA, showing the relationship between gene expression and samples*

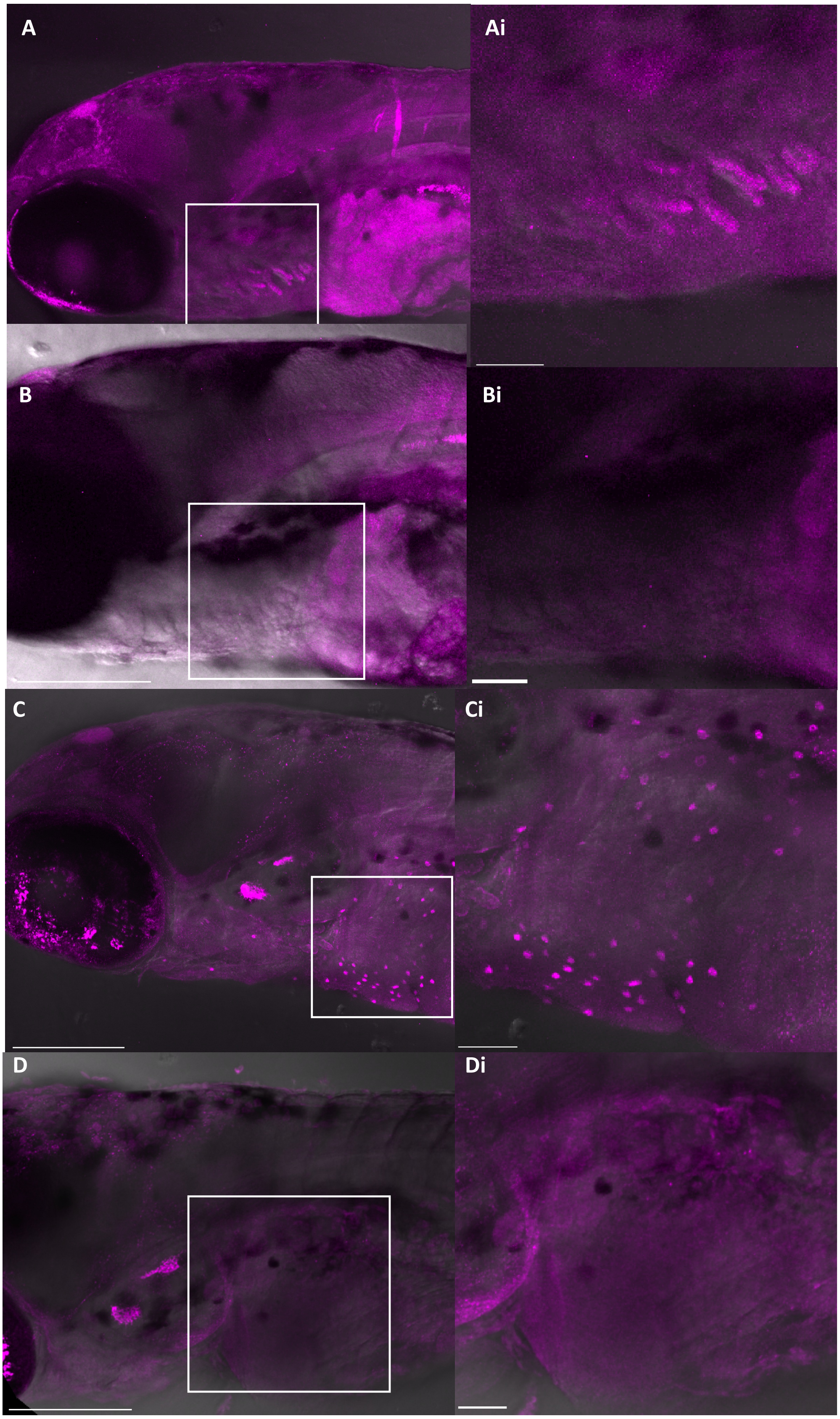


***Fig. S4: Confocal imaging of rhesus glycoproteins in F0 crispant 5 dpf zebrafish.*** *A) Rhbg antibody (1:250) in Rhcgb-crispant zebrafish. A brightfield image is used for reference and fluorescent signal of Rhbg-antibody (magenta) is shown. Ai is a detail of the white square area showing signal of Rhbg-antibody on the developing gill filaments. B) Rhbg antibody (1:250) in Rhbg-crispant zebrafish. A brightfield image is used for reference and*

*fluorescent signal of Rhbg-antibody (magenta) is shown. Bi is a detailed image of the white square area showing the absence of signal of Rhbg-antibody on the developing gill filaments as in Ai. C) Rhcgb antibody (1:500) in Rhbg-crispant zebrafish. A brightfield image is used for reference and fluorescent signal of Rhcgb-antibody (magenta) is shown. Ci is a detail of the white square area showing signal of Rhcgb-antibody on the yolk sac ionocytes. D) Rhcgb antibody (1:500) in Rhcgb-crispant zebrafish. A brightfield image is used for reference and fluorescent signal of Rhcgb-antibody (magenta) is shown. Di is a detail of the white square area showing the absence of Rhcgb-antibody on the yolk sac ionocytes. For all overview images, the scale bar indicates 200 µm, for all inserts the bar indicates 50µm.*

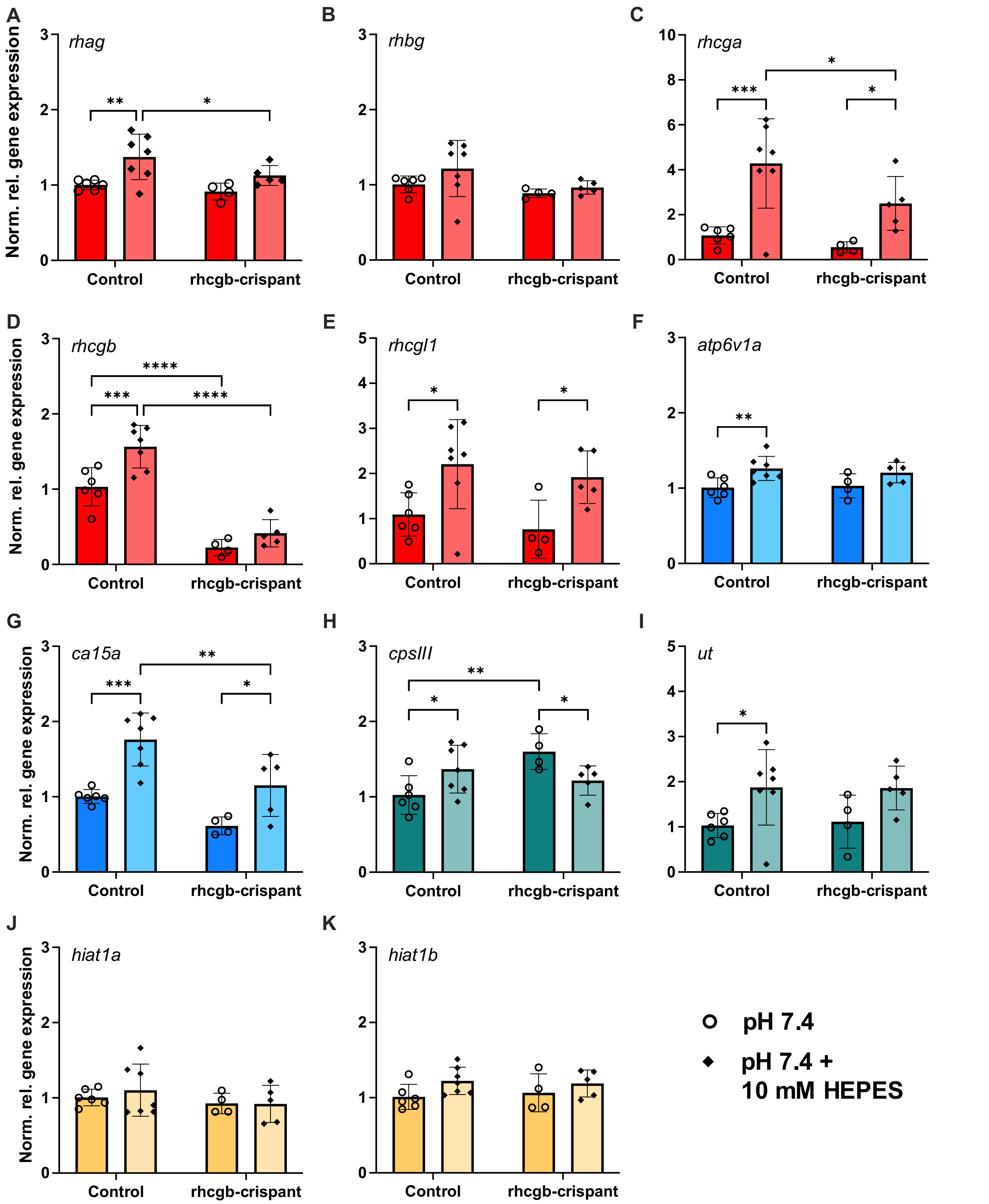


***Fig. S5: Effect of HEPES-buffering on nitrogen metabolic and ionocyte-related gene expression in control and rhcgb-crispant zebrafish.*** *Control zebrafish and crispants were exposed to 10 mM HEPES (pH 7.4) for 3 hours and whole-body expression levels of nitrogen metabolic genes and ionocyte-markers were measured. Expression of rhesus glycoproteins (rhag, rhbg, rhcga, rhcgb and rhcgl1;* ***A-E****), HR-ionocyte markers (atp6v1aa, ca15a;* ***F-G****). urea metabolism genes (cpsIII, ut (****H-I****) and hippocampal-abundant transcripts 1a and 1b (hiat1a, hiat1b;* ***J-K****) was measured in pooled embryonal and larval samples (n = 4-7 per group). Gene expression is expressed as normalized relative gene expression (normalized to unexposed control fish samples and calculated relative to the expression of two housekeeping genes: β-actin and 40S ribosomal protein 11 (RPS11). Individual values are shown*

for each group, with a colored bar and error bars indicating mean relative expression ± SD. Bar colors correspond to gene categories (red: rhesus glycoproteins, cyan: ionocyte-marker, yellow: hiat-ammonia transporters, teal: urea metabolism). Unexposed zebrafish reared at pH 7.4 are shown as open circles, while zebrafish exposed to HEPES are shown as closed diamonds. Significance between crispants and control fish and between HEPES-exposed fish and control conditions was tested with a two-way ANOVA and Tukey post-hoc tests. Asterisks indicate the significance level of Tukey post-hoc tests (* $p < 0.05$, ** $p <0.01$), *** $p < 0.001$, **** $p < 0.0001$).
